# In-feed bacitracin methylene disalicylate alters microbiota function and increases antibiotic resistance in a dose-dependent manner

**DOI:** 10.1101/2025.02.16.638580

**Authors:** Paul Oladele, Carmen L. Wickware, Julian Trachsel, Torey Looft, Timothy A. Johnson

## Abstract

Antibiotics are commonly used in turkey production to prevent and treat infection which can improve animal growth and efficiency, but the mechanism by which antibiotics improve animal performance, and the resistance risks associated with the antibiotic inclusion levels remain unclear, particularly in turkey production. Therefore, we investigated the longitudinal effect of subtherapeutic and therapeutic doses of bacitracin methylene disalicylate (BMD) on antibiotic resistance genes, mobile genetic elements, and metabolism genes by analyzing the turkey cecal metagenome. The therapeutic dose of BMD increased a vast array of antibiotic resistant genes (ARGs), conjugation-related genes of type IV secretion system and transduction-related genes for the length of the experiment (78 days), while a smaller, transient effect was observed due to the subtherapeutic dose. Estimated bacterial growth rate, estimated by metagenome assembled genome sequence coverage, decreased after 7 days of in-feed BMD, but increased in the therapeutic group over time. Tryptophan synthesis from chorismate increased in a dose-dependent manner between days 7 - 35. Overall, the effects of subtherapeutic BMD on the turkey cecal microbiota was temporary while those of a therapeutic dose were longer lasting. This study shows that antimicrobial resistance genes belonging to multiple antibiotic classes, and mobile genetic elements (MGEs) were increased after BMD administration. The enrichment these genes by BMD shows the risk associated with antimicrobial feed additives. BMD’s effect on tryptophan synthesis provides a potential metabolic target for developing non-antibiotic microbiome modulatory growth promoters for turkey production.

**Importance:** Antibiotic use in agricultural animals remains a hotly debated and important topic to human, animal, and environmental health. The dose-dependent responses to BMD, an antibiotic fed additive allowed at both therapeutic and subtherapeutic doses, are not well understood. This study highlights that therapeutic use of BMD is a stronger selective pressure than the subtherapeutic dose for antibiotic resistance genes and genes related to horizontal gene transfer. This indicates that use of BMD could select for antibiotic resistant bacteria that may pose a risk to animal, human and environmental health. Additionally, this study highlighted that BMD decreased the activity of beneficially bacteria, and therefore may be associated with a decreased concentration of bacterial metabolites (especially tryptophan related) in the cecum. This may indicate that growth promoting antibiotics suppress bacterial activity generally, rather than allowing beneficial bacteria to generate beneficial metabolites.

## Introduction

In livestock production, antibiotics have been widely used to treat infection and promote animal growth (Chattopadhyay, 2014; Damron et al., 1991; Dibner & Richards, 2005). Both subtherapeutic (Feighner & Dashkevicz, 1987) and therapeutic (Gupta et al., 2021) doses of antibiotics have been shown to improve animal growth and feed efficiency. While therapeutic antibiotics are prescribed in specific cases of bacterial infection, both dosage levels contribute to the increasing incidence and spread of antibiotic resistance genes, posing a significant threat to public health (Aslam et al., 2018; Gaskins et al., 2002; Landers et al., 2012; Manyi-Loh et al., 2018).

In response to this growing public health concern regarding antibiotic resistance in animal agriculture, legislation against the use of antibiotic growth promoters (AGPs) has increased. For instance, the European union banned all AGPs in 2006, and US Food and Drug Administration (FDA) implemented the veterinary feed directive (VFD) in 2017 which mandates veterinarian oversight when medically important antibiotics are administered in animal feed (FDA, 2013). This led to a reduction in overall antibiotic use in livestock production. For example, the Netherlands achieved a 70% reduction in antibiotic use in animals by 2016 compared to 2009 levels (Dorado-García et al., 2016) and in the United States, antibiotics used in animal agriculture dropped from 13.98 million kilograms in 2016 to an average of 11.11 million kilograms between 2017 – 2020 (FDA, 2021). However, the reduction was not uniform across all sectors. Antibiotic use in chicken production declined 30%, from 2.18 million kg in 2016 to 1.52 million kg between 2017 and 2020, whereas in turkey production, it decreased 15%, from 1.15 million kg to 0.98 million kg during the same period (USDA *ERS*, 2022). Given the extensive use of antibiotics in turkey production, developing alternatives and assessing antibiotic resistance risks are crucial (Wallinga et al., 2022).

Despite the high use of antibiotics, the mechanisms by which it enhances performance and their impact on the microbiome remain poorly understood. Antibiotics may improve performance by modulating the microbiome, either by reducing gut pathogen density (Chattopadhyay, 2014) or altering the taxonomic and functional composition of gut microbiota (Díaz Carrasco et al., 2018; Proctor & Phillips, 2019) and metabolome (Plata et al., 2022) to benefit host health (Dibner & Richards, 2005). Understanding AGP-induced changes in the gut microbiome composition and function is crucial, as the microbiome can influence host health directly via physical contact or indirectly through fermentation metabolites that modulate immune development and intestinal barrier function.

In addition to growth promotion, in-feed antibiotics increase the abundance of antibiotic-resistant bacteria. Studies have shown that subtherapeutic and therapeutic antibiotics affected the resistome of chicken microbiota differently (Gupta et al., 2021). Subtherapeutic antibiotics have been thought to increase selection for resistance due to prolonged exposure compared to shorter use at therapeutic levels (Zaheer et al., 2013). In chickens, subtherapeutic BMD has elevated bacitracin resistance genes, while therapeutic dose increased multidrug resistant isolates (Gupta et al., 2021; Zhang et al., 2024). Keijser et al. (2019) observed a dose-dependent rise in the resistome of veal calves after oxytetracycline administration, and Zaheer et al. (2013) reported an increase in resistant enterococcus isolated from beef cattle treated with either therapeutic or subtherapeutic macrolides. These findings suggest that selection pressure for ARGs is dependent on antibiotic concentration.

Compared to chicken and cattle, limited studies have investigated the impact of antibiotics on the functional and resistome profiles of the turkey microbiome, despite the historically high use of antibiotics in turkeys. Understanding how subtherapeutic and therapeutic doses influence these profiles could improve our understanding of the risks associated with different antibiotic dosage and the mechanism by which AGPs enhance performance. This knowledge could help uncover new hypotheses that may lead to the development of non-antimicrobial growth promoters without increasing selection for antibiotic resistance genes.

We previously showed that both therapeutic and subtherapeutic doses of in-feed bacitracin methylene disalicylate (BMD) – a polypeptide AGP that inhibits bacterial cell wall synthesis and is allowed in animal feed at both dosage in the United States (FDA, 2023) – reduced microbiome richness and altered the cecal metabolome in turkey (Johnson et al., 2019). BMD administration initially (day 7) decreased the concentration of many metabolites, followed by a general increase in metabolites until day 87. BMD also increased the concentration of tryptophan and altered the concentration of many tryptophan metabolites in cecal contents of turkeys. However, it remains unclear whether these changes were driven by shifts in bacterial metabolism or nutrient uptake by the host. The impact of BMD on the antibiotic resistome is also not yet fully understood.

Our objectives were to determine the effect of therapeutic and subtherapeutic doses of BMD on the functional characteristics and antimicrobial resistance gene content in turkey cecal microbiota. We hypothesized that both subtherapeutic and therapeutic doses of BMD would reduce bacterial metabolic potential, alter aromatic amino acid metabolism, and increase antibiotic resistance genes in the turkey cecal microbiota.

## Results

Shotgun metagenomic sequencing of cecal content from 90 turkeys at 7, 35 and 78 days after experimental diet initiation generated 2,480,187,759 sequences from Illumina HiSeq sequencing and 7,178,582 sequences from PacBio sequencing. After sample processing and quality control, reads were co-assembled and 594,500 open reading frames (ORFs) were identified and annotated using the TIGRfam database. To confirm the absence of bias in annotation rate by BMD treatment or age, we correlated the total number of sequences with the number of TIGRfam annotations, yielding a strong correlation (*r^2^ =* 0.95, *p >* 0.0001) across all ages and treatment group, without discernible bias (Fig. 1a).

**Figure 1.**
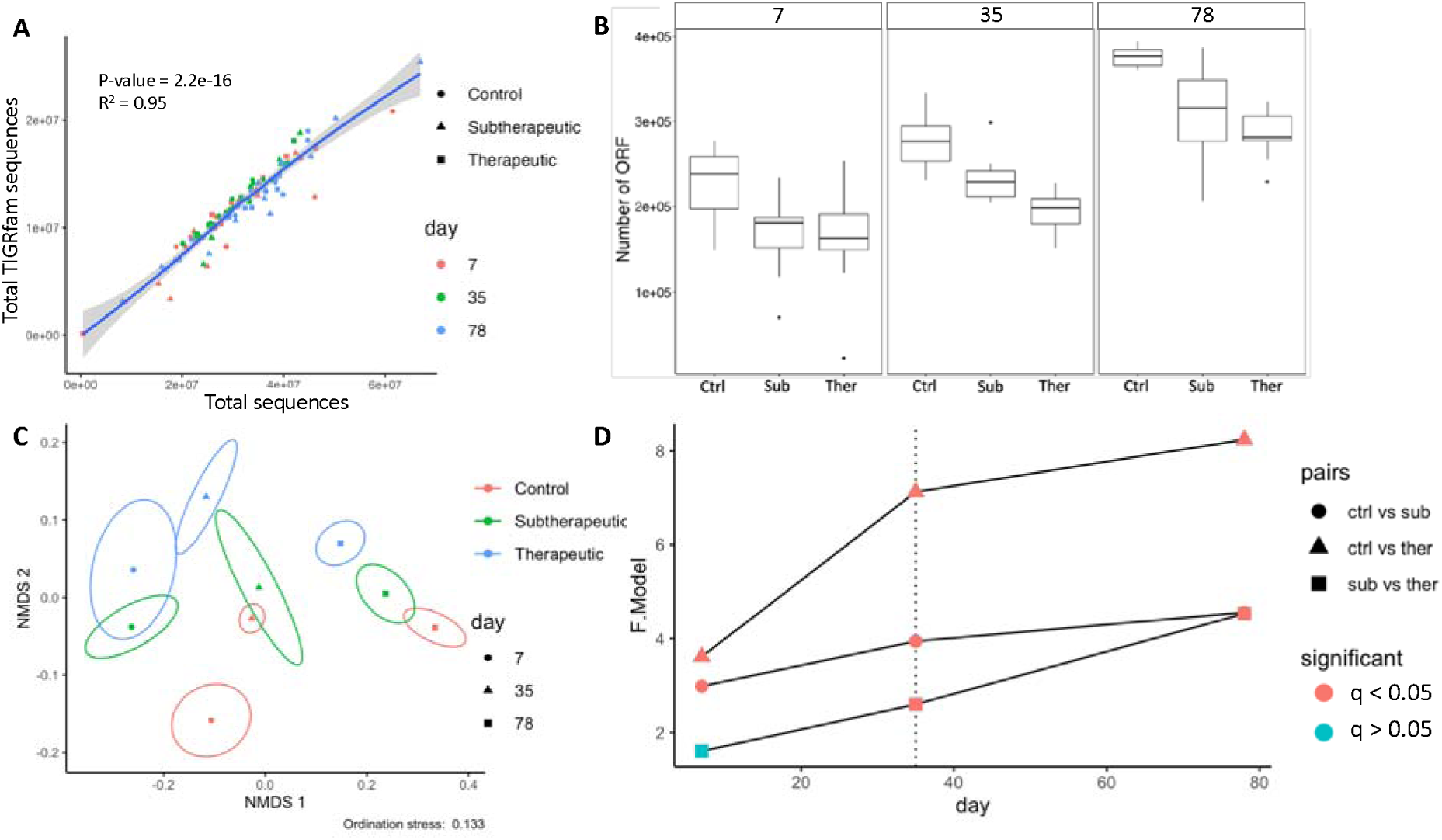
Diversity metrics of ORFs. A. Correlation plot of TIGRfam annotation and Total sequences. B. Number of ORFs in each treatment groups within each day. C. Non-metric multidimensional scaling (NMDS) ordination (Bray-Curtis) of ORFs showing the effect of both BMD administration and age with stress shown below the figure. Ellipses indicate 95% confidence interval with the symbol at the centroid of all replicate samples of BMD treatment (indicated by color) and age (indicated by shape). D. PERMANOVA test for differences in community structure due to BMD treatment and age. F-value indicate the degree of difference between two communities compared, shapes indicate pairwise BMD treatment comparison and color indicate pairwise comparison statistical significance. P-value was adjusted for false discovery rate method for multiple comparisons.

### BMD treatment decreased ORF diversity and shifted functional profile

The number of observed ORFs decreased dose-dependently with BMD treatment, with the therapeutic dose having the fewest number of ORFs (*p* < 0.05). ORF numbers decreased in all BMD-treated groups but increased with age (Fig. 1b). We further assessed the impact of BMD and age on the functional gene profile of the turkey cecal microbiota using non-metric multidimensional scaling (NMDS) based on Bray-Curtis dissimilarity. Both BMD treatment and age significantly influenced the structure of the ORFs in the cecal microbiome (PERMANOVA, *q <* 0.05). The diversity of ORFs shifted along the y-axis in a BMD dose-dependent manner and along the x-axis with age (Fig. 1c). Interestingly, the therapeutic group remained distinct from the subtherapeutic group through day 78, despite receiving the same BMD dose as the subtherapeutic group from day 35 onward (PERMANOVA, *q <* 0.05, Fig. 1c). Compared to the control group, the therapeutic group showed a greater magnitude of beta diversity shift than the subtherapeutic group, while the rate of change decreased after day 35 (Fig. 1d). This suggests that a therapeutic dose of BMD had a greater impact on the functional profile of the turkey cecal microbiota than the subtherapeutic dose.

### BMD treatment impacted the abundance of functional genes

In addition to the global effects of BMD on the functional gene profile, its impact on various functional pathways was investigated by performing differential abundance analysis using DESeq2. Open reading frames (ORFs) were grouped into 2561 gene functions based on TIGRfam annotation for analysis. Compared to the control group, the subtherapeutic group had 292, 235 and 234 differentially abundant genes (increased: 115, 144, 135; decreased: 177, 91, 99) on days 7, 35 and 78, respectively. In the therapeutic group, 538, 1,044, and 1,393 genes were differentially abundant (increased: 282, 562, 732; decreased: 256, 482, 661) on days 7, 35 and 78, respectively (Fig 2a; Table S1 - 6). Compared to the control, the therapeutic group consistently had more differentially abundant genes than the subtherapeutic group both in terms of upregulated and downregulated genes. This suggests a stronger impact of the therapeutic BMD on the microbiota functional gene profile.

**Figure 2.**
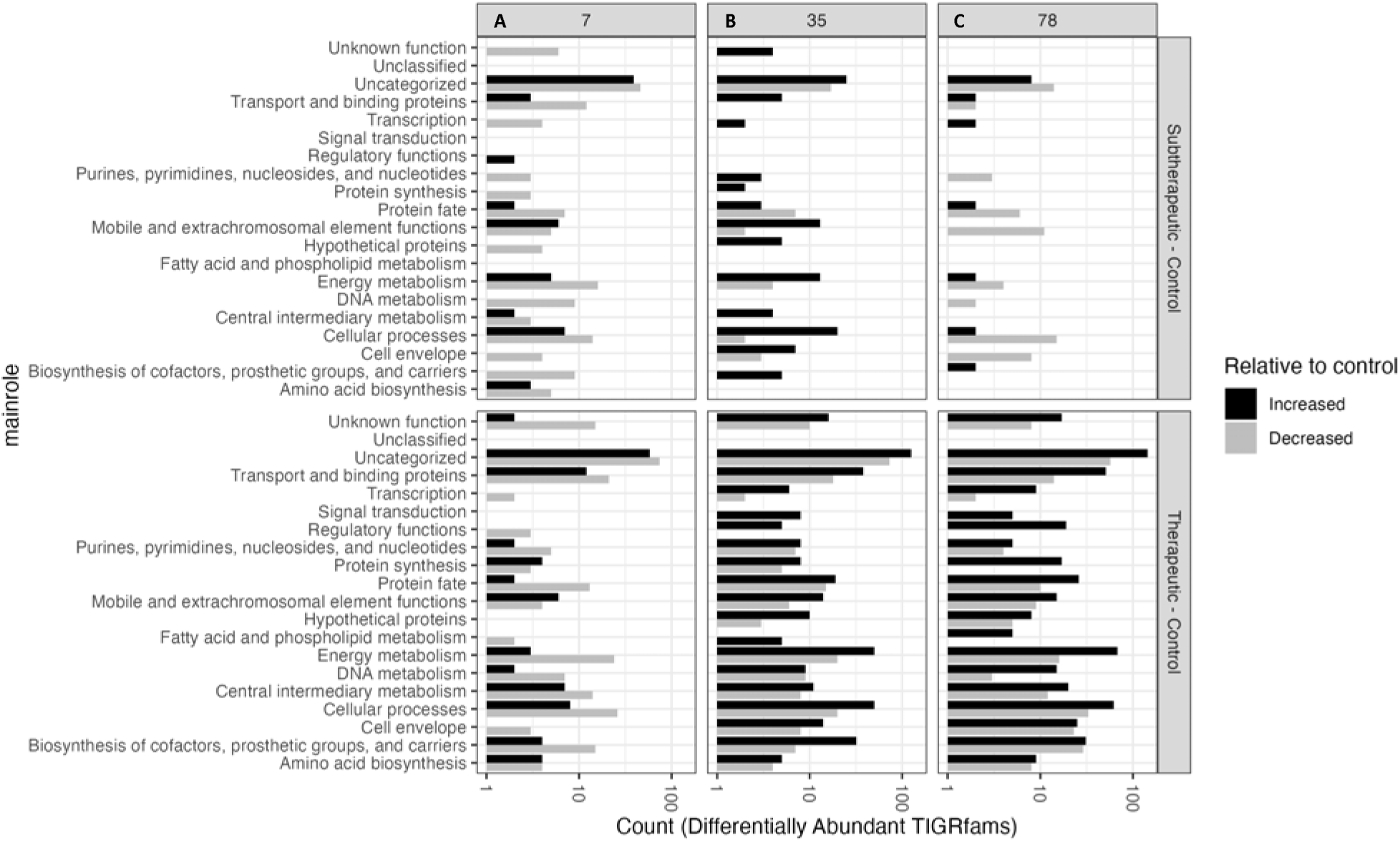
Count of differentially abundant genes from DESeq2 in treatment groups relative to control. A - C Subtherapeutic (upper) and Therapeutic (lower) dose on days 7, 35 and 78, respectively. Statistical significance was determined with DESeq2. Black color indicates DAGs that are significantly increased while grey indicates DAG that are significantly decreased (q<0.05).

### BMD treatment impacted resistome profile of cecal microbiome of turkey

Previous studies have demonstrated that antibiotics impact the profile of antimicrobial resistance genes in the both human (reviewed in Schwartz et al., 2020) and animal (reviewed in Xu et al., 2022) microbiomes. To investigate how BMD affects the cecal resistome in turkeys, we identified 393 antibiotic resistance genes (ARGs) across the samples using the Resistance Gene Identifier (RGI, version 4.0.3) based on the Comprehensive Antibiotic Resistance Database (CARD, version 3.2.0) (McArthur et al., 2013). The profile of ARGs in the turkey cecum was impacted by both antibiotic treatment and age (Fig. 3a), while BMD caused a dose-dependent shift in ARG profiles. On day 7, no significant difference was observed between the subtherapeutic and therapeutic group (PERMANOVA, *q >* 0.05), but both groups differed significantly from the control (PERMANOVA, *q <* 0.01). By days 35 and 78, both the subtherapeutic and therapeutic group were distinct from the control group (PERMANOVA, *q <* 0.05), as well as from each other (PERMANOVA, *q <* 0.05). ARG diversity also shifted over time, similar to the pattern seen with functional genes. The ARG community structure dissimilarity between the therapeutic and control groups increased steadily through day 78. In contrast, the ARG community structure dissimilarity between the subtherapeutic and control groups decreased after day 7 (Fig. 3b). This suggests that therapeutic BMD may have a longer-lasting impact on the resistome.

**Figure 3.**
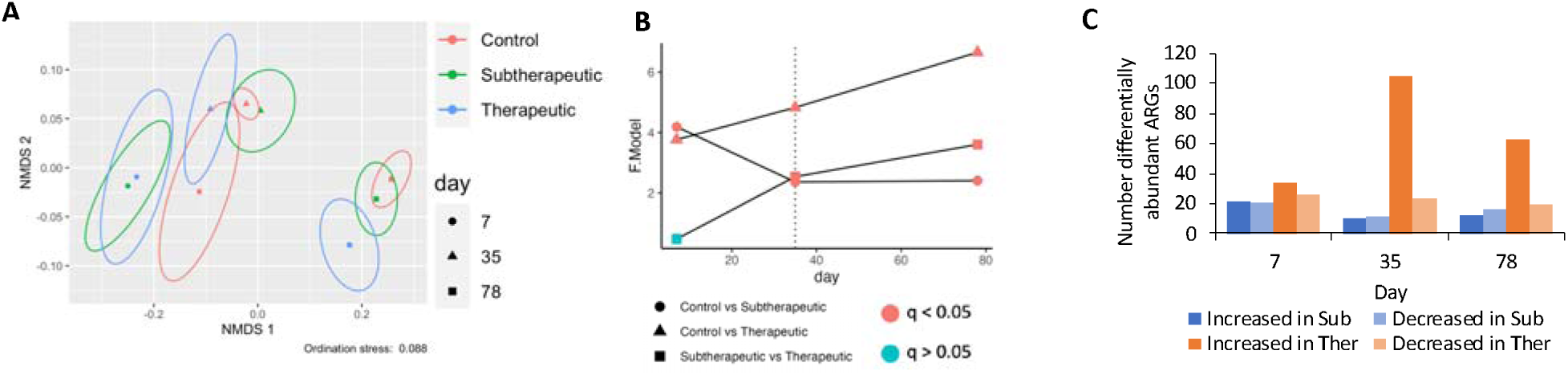
A. Non-metric multidimensional scaling (NMDS) ordination (Bray-Curtis) of ARGs showing the effect of both BMD administration and age with stress shown below the figure. Ellipses indicate 95% confidence interval with the symbol at the centroid of all replicate samples of BMD treatment (indicated by color) and age (indicated by shape). B. PERMANOVA test for differences in community structure due to BMD treatment and age. F.Model indicates the degree of difference between two communities compared, shapes indicate pairwise BMD treatment comparison and color indicate pairwise comparison statistical significance. P-value was adjusted for false discovery rate method for multiple comparisons. C. Number of differential abundant ARGs on day 7, 35 and 78 in all treatment groups.

To further explore the changes in the resistome structure, we performed a differential abundance analysis using DESeq2 to identify ARGs whose abundance was altered by BMD. The therapeutic group exhibited 3-5 times more differentially abundant ARGs (relative to the control) on days 35 and 78 than the subtherapeutic group (Fig. 3c, detailed results in Table S10 - 15). On day 7, both treatment groups had a similar number of increased and decreased ARGs (Fig. 3c and Table S7 - 9).

Over time, BMD administration affected a wide variety of ARGs, but antibiotic efflux and antibiotic inactivation were the dominant ARG mechanisms in both treatment groups (Fig. 4). When turkeys were given a subtherapeutic dose, bacitracin resistance gene (*bacA*) increased only on day 7 but not later. In contrast, *bacA* increased across all days in the therapeutic dose (Fig. 4a - c). The two-component regulatory system (*baeR* and *baeS*), which is involved in bacitracin resistance, also increased only in the therapeutic group. Despite the reduction of BMD to subtherapeutic levels on day 78, the therapeutic group still exhibited elevated ARG levels. Additionally, ARGs conferring resistance to tetracycline (*tet*(44), tet(45), *tet*(L), *tet*(W/N/W), *tet*(Z), *tet32*, *tetA*(P), *tetB*(P), *tetM*, *tetO* and *tetS*), vancomycin (a D-alanine-D-serine ligase vanG cluster, labelled as vanI), and aminoglycosides (*aph*(6)-Id, *aph*(3’’)-Ib and *aph*(3’)-IIIa) were increased in the therapeutic group (Fig. 4b - c) showing that the increased ARGs were not limited to bacitracin resistance. Multidrug efflux pump genes (*acrB*,*D*,*E*,*F*, *cpxA*, *emrA*, *emrK*, *evgS*, *gadX*, and *mdtA,B*,*C*,*E*,*F*,*G,H,M,N,O,P*) increased primarily in the therapeutic group (Fig. 4b - c).

**Figure 4.**
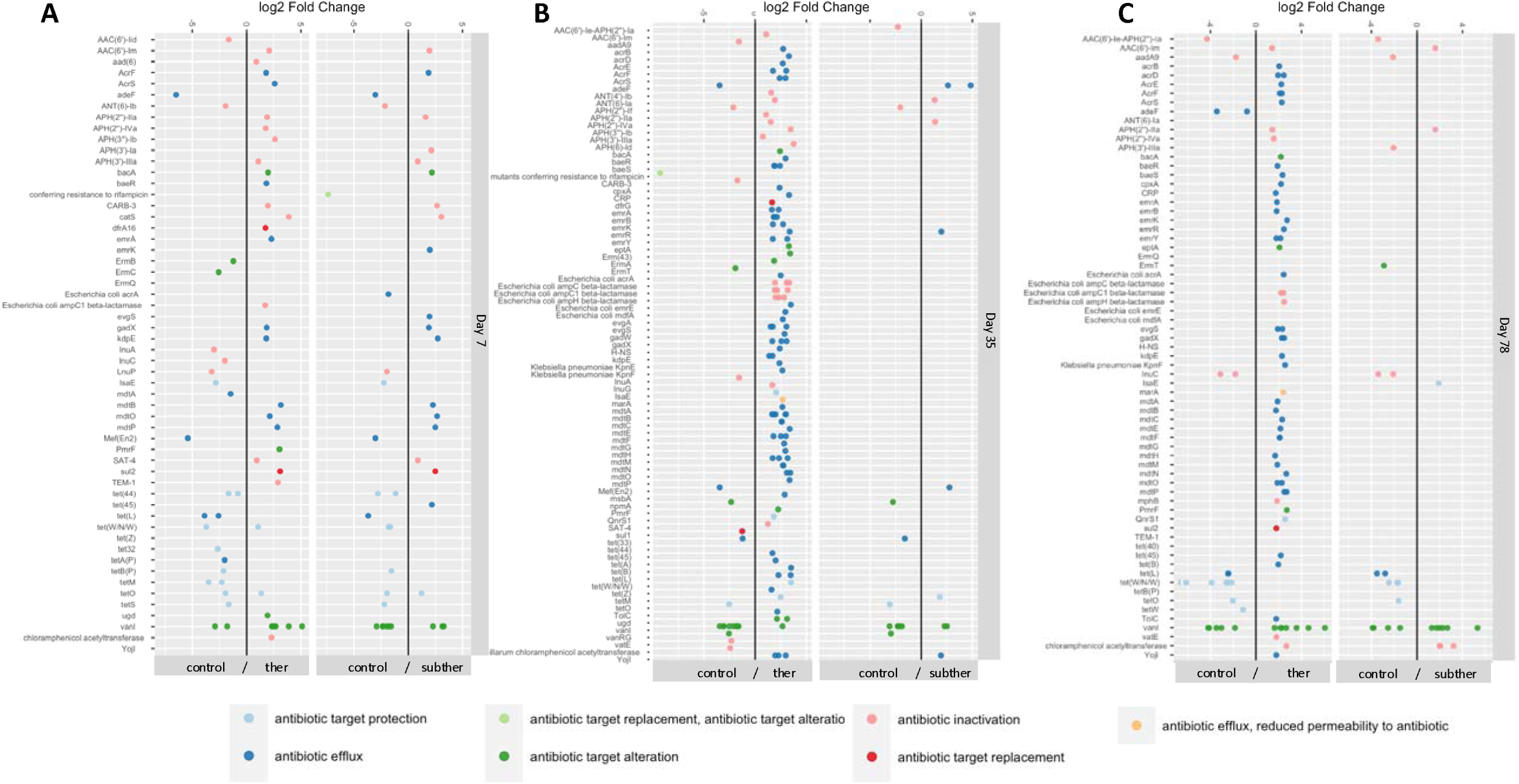
Differential abundant ARGs using DESeq2 between both therapeutic and control, and subtherapeutic and control on A. day 7, B. day 35 and C. day 78.

### BMD increased the abundance of mobile genetic elements

To access the impact of BMD on specific cellular functions, we compared differentially abundant genes (DAGs) in various metabolic pathways. On day 7, in both the subtherapeutic and therapeutic groups, 15 gene categories had more DAGs that were decreased (mostly related to essential cellular functions), while only one category had more DAG that were increased (mobile and extrachromosomal element functions) (Fig. 2a; Fig. S1a). By day 35 and 78, in the therapeutic group, the pattern reversed. All metabolic gene categories had more DAGs that were increased compared to control group (Fig. 2b; Fig. S2 and 3). In the subtherapeutic group on day 35, most categories had more DAGs that were increased compared to the control, but this reversed on day 78 when energy, nucleotide and central metabolism all had more decreased DAGs. There were more DAGs related to mobile and extrachromosomal functions that increased in abundance than those that decreased in abundance on all sampling days in the therapeutic group, while in the subtherapeutic group, more MGEs were increased than decreased only on days 7 and 35.

### BMD increased the abundance of F-like bacterial conjugation genes and phage related genes

Given the increase in mobile genetic elements in both the subtherapeutic and therapeutic groups, we aimed to identify which specific genes were impacted. We found two key types of mobile genetic elements whose abundance was altered by BMD treatment: type IV secretion system (T4SS) used for conjugation and phage-related genes involved in transduction. Since some of the TIGRfam genes were not categorized, we expanded our search to include genes in the uncategorized category that could be identified as related to mobile genetic elements. In both the subtherapeutic and therapeutic groups, about 20 T4SS genes increased in abundance on day 7 (Fig. 5a and b). On day 35, the subtherapeutic group showed an increase in only 2 T4SS genes, while 5 were increased in therapeutic (Fig. S4a and b). On day 78, T4SS differential abundance was restored in the therapeutic group (14 DAG increased, Fig. 5d) but not in subtherapeutic group (0 DAG, Fig 5c). In both BMD treatment groups, 5 other MGE genes associated which conjugative transposons either decreased due to BMD or were not differentially abundant (Fig. S5).

**Figure 5.**
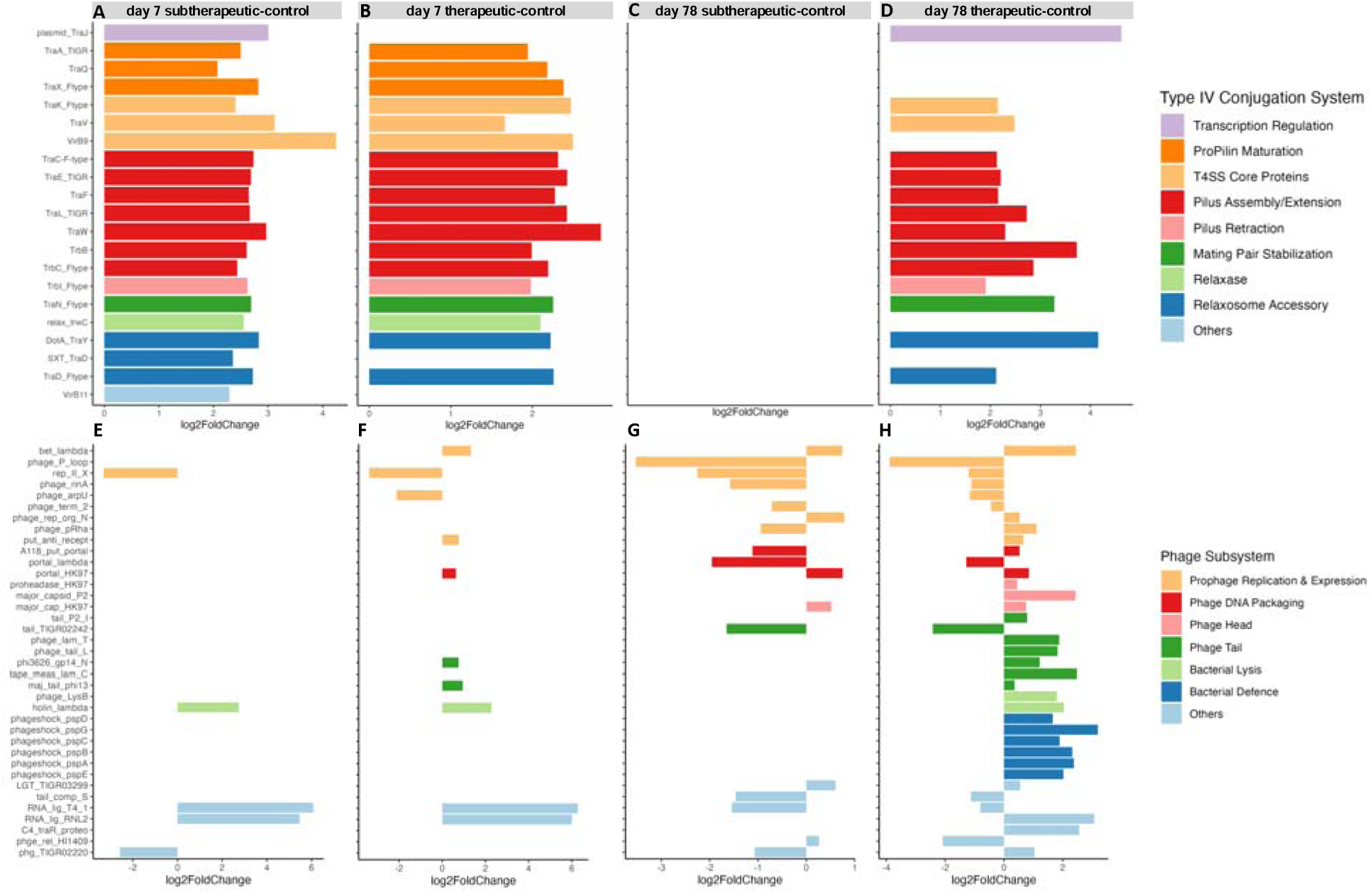
Differentially abundant genes of the type IV secretion system (T4SS) use for conjugation relative to control on day 7 in A. Subtherapeutic, B. Therapeutic, on day 78 in C. Subtherapeutic and D. Therapeutic. Differentially abundant phage related genes relative to control on day 7 in E. Subtherapeutic, F. Therapeutic, on day 78 in G. Subtherapeutic and H. Therapeutic.

We also investigated prophage-related genes. Most prophage-related TIGRfams were classified in the mobile elements category (34/42 TIGRfams). BMD administration led to differential abundance of these phage-related genes in both treatment groups. On day 7, no major change in phage-related gene abundance was observed in either group (Fig 5e and f). On day 35, 18 genes increased in abundance in both BMD groups, while 2 and 5 genes decreased in abundance in the subtherapeutic and therapeutic groups, respectively (Fig. S4c and d). On day 78, 6 genes increased, and 11 genes decreased in abundance in the subtherapeutic group compared to the control group, while and 27 genes increased, and 10 genes decreased in abundance in the therapeutic group (Fig 5g and h). The phage genes were associated with steps in prophage induction, including prophage replication, phage DNA packaging, phage head, phage tail, bacterial lysis, and bacterial defense against phage.

### Co-occurrence analysis of conjugation and phage genes with antibiotic resistance genes

Given the observed increase in F-like T4SS genes and phage-related genes (MGEs), which coincided with increase in ARG abundance, particularly in the therapeutic group on day 78, we conducted a co-occurrence analysis (Spearman’s correlation, *rho* = 0.8, *q* < 0.05) to assess the association between ARGs and the MGEs. This analysis aimed to investigate the potential mobility of ARGs on MGEs. ARG-MGE co-occurrence networks were constructed with either conjugation genes or phage genes. Both T4SS and phage genes showed significant co-occurrence (*q* < 0.05) with ARGs, but the complexity of the network changed between the timepoints (Table S3).

On day 7, the control and subtherapeutic group ARG-conjugation gene co-occurrence networks had higher numbers of edges than the therapeutic group (Fig. S6, Table S16). However, both BMD treatments generated phage networks with less complexity than the control group (Fig. S8). On day 35, network structures had low complexity across all treatment groups for both T4SS (Fig. S7) and phage networks (Fig. S9). By day 78, the therapeutic group had the highest number of ARG-MGE interactions of all treatment groups (Fig. 6). A similar pattern was observed for the relatively simple ARG-phage gene co-occurrence on day 78, in which the therapeutic group had the highest network complexity (Fig. S10, Table S16). Taken together, there was a decrease in network complexity in each treatment group over time, except that the therapeutic group had its most complex ARG-T4SS network on day 78 and a slightly more complex ARG-phage network on day 78 than on day 35. On day 78 in the therapeutic group, *traC* (degree = 22) had the highest degree in the ARG-T4SS network (Fig. 6c and Table S16) while phage tail assembly protein T (degree = 25) had the highest degree in the ARG-phage network (Fig. S10c and Table S16).

**Figure 6.**
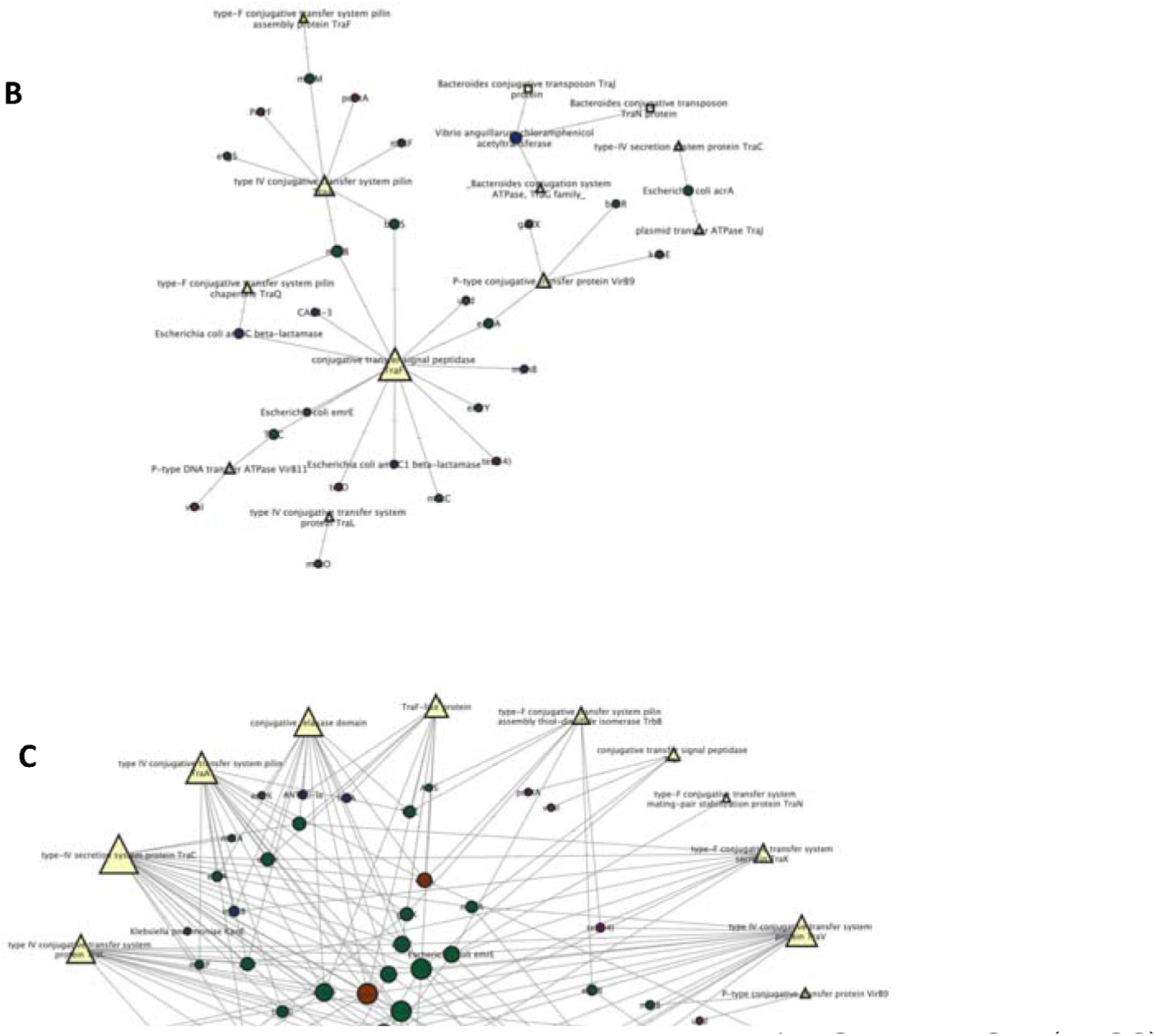
Network analysis showing the spearman correlation between ARG and MGE (T4SS) in turkey cecal microbiota on day 78. Pairwise correlation of padj < 0.05 and rho > 0.8 was considered. The nodes represent the ARG and MGE, the size of the node represent its number of connections (Degree). A-C represent ARG– conjugation gene (both T4SS and conjugative transposon) on day 78 in control, subtherapeutic and therapeutic group. Triangles are T4SS, square are conjugative transposon while circle are ARG. Color of the circle indicate different ARG mechanism, green is antibiotic efflux, blue is antibiotic inactivation, red is antibiotic target alteration, purple is antibiotic target protection while light green is antibiotic target replacement.

### BMD enriched genes involved in tryptophan biosynthesis

Based on our previous observation of that tryptophan metabolites were altered by BMD (Johnson et al., 2019), we conducted further analysis of the genes within the amino acid synthesis pathway, which revealed that genes involved in aromatic amino acid metabolism, particularly in tryptophan biosynthesis from the shikimate pathway and phenylalanine through chorismate were changed by BMD treatment. Tryptophan synthase alpha and beta chains, which are responsible for the conversion of indole and (3-Indolyl)-glycerol phosphate to tryptophan, were increased by both doses of BMD on day 7 and 35. Additionally, genes involved in the conversion of phenylalanine to tryptophan through chorismate with anthranilate as an intermediate, and the conversion of quinate to tryptophan through chorismate with shikimate as intermediate, were also increased by BMD in a dose-dependent manner (Fig. 7a-h, Fig. S11).

**Figure 7.**
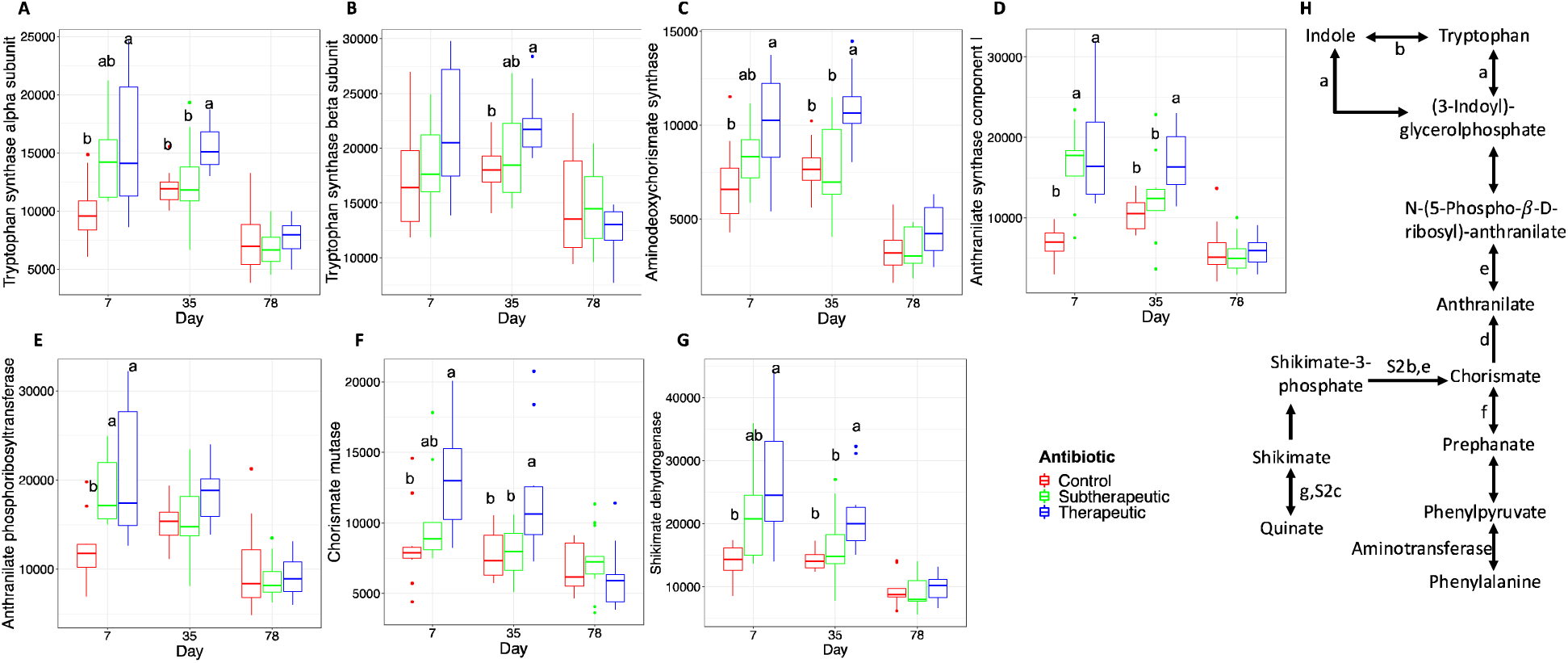
Abundance of genes involved in phenylalanine synthesis from either tryptophan or quinate. A. Tryptophan synthase alpha subunit. B. Tryptophan synthase beta subunit. C. Aminodeoxychorismate synthase. D. Anthranilate synthase component I. E. Anthranilate phosphoribosyltransferase. F. Chorismate mutase. G. Shikimate dehydrogenase. H. Schematic representation of the pathway for reference.

### Estimated growth rate of bacteria decreased with therapeutic administration of BMD but recovered over time

Bacterial growth rates were estimated using the index of replication (iRep), which measures the genome sequencing coverage across the genome and is quantified by differential sequencing coverage at origin (peak) and terminus (trough) of replication. We obtained valid iRep estimates from 161 metagenome-assembled genomes (MAGs). Both therapeutic and subtherapeutic BMD treatments were associated with reduced community-average iRep growth rate estimates compared to the control group. The largest reductions in iRep estimates occurred on day 7, with the therapeutic group showing a difference of −0.20 (P < 0.001), while the subtherapeutic group showed a difference of −0.07 (P = 0.003) (Fig. 8a, and b; Fig. S12a and d). Several MAGs, such as bin231 (*Faecalibacterium prausnitzii*), bin235 (*Coprococcus comes*), bin280 (*Dorea formicigenerans*), bin433(*Anaerostipes caccae*), bin886 (*Erysipelotrichaceae bacterium*) and bin493 (f_Lachnospiraceae) showed significant reduction in iRep due to BMD treatments, particularly in the therapeutic group. On day 35, there was no overall difference in iRep estimates. On day 78, small but significant overall reductions in iRep growth rates were associated with both the therapeutic (difference = −0.07, P= 0.006), and subtherapeutic (difference = −0.06, P = 0.01) treatments compared to controls (Fig. 8a and b; Fig. S12c and f).

**Figure 8.**
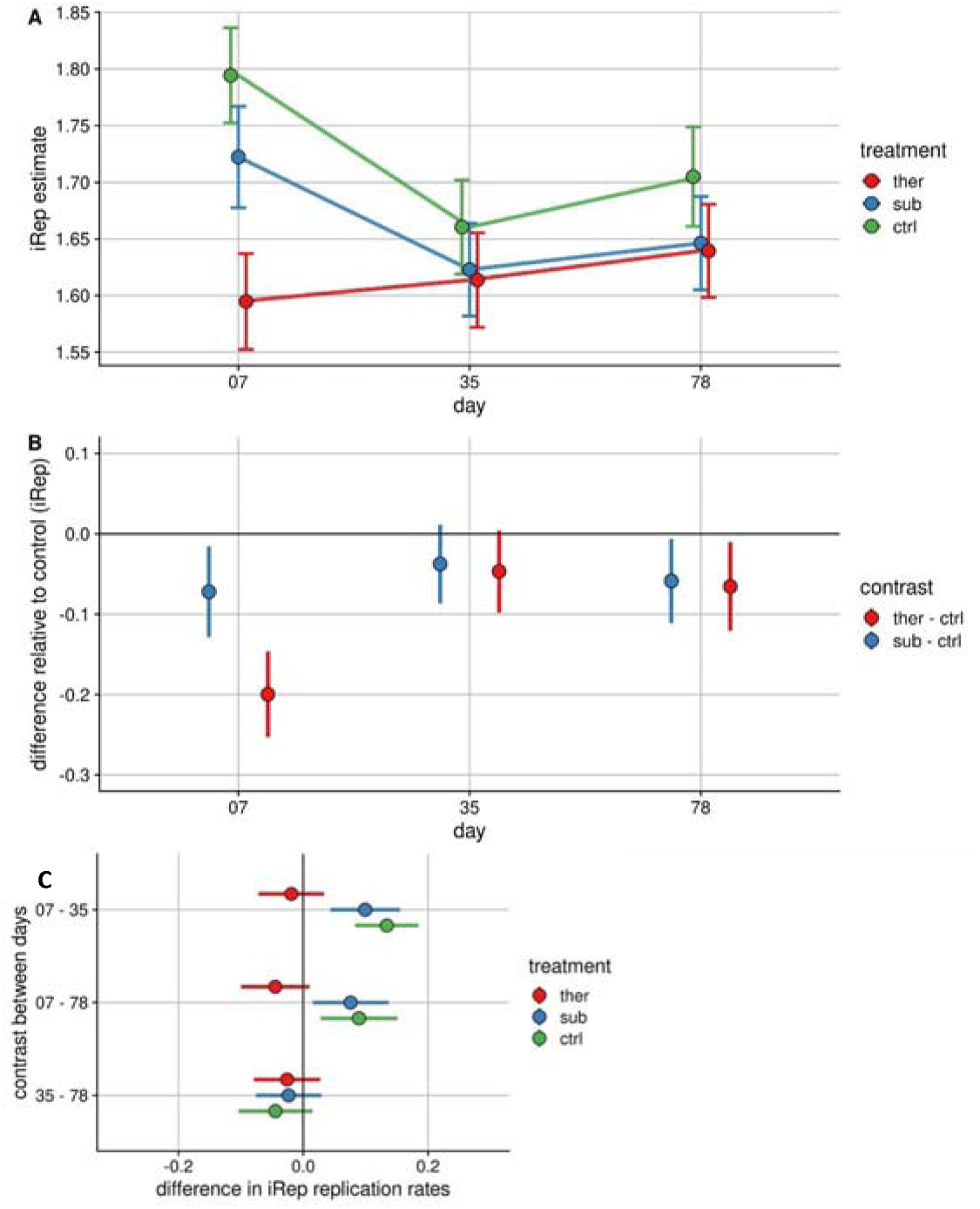
Effect of BMD administration on community wide replication rates across time. Estimated growth rate of bacteria decreased with therapeutic administration of BMD but recovered over time. Effect of BMD administration on community wide replication rates across time. **A.** Estimated marginal means for iRep replication rates with 95% confidence intervals around the estimates at each day for each treatment. **B.** Treatment contrasts at each day depicting the difference in iRep replication rates between the control and each experimental group at each day. The estimated difference is represented by the points with 95% confidence intervals around those estimates. **C.** Differences in community wide replication rates over time in control and BMD fed animals. Each point represents the difference in iRep replication rates between the indicated days within each treatment. The lines represent 95% confidence intervals around the estimated differences. iRep estimates from MAGs with at least 4 observations in all treatment groups at each timepoint were included in the analysis.

## Discussion

Antibiotic use in U.S. turkey production, relative to the quantity of meat produced, remains relatively high compared to other livestock species (Wallinga et al., 2022), but our understanding of its impact on the risk of antibiotic resistance is currently limited. To address this gap, we examined the effects of BMD - an antibiotic approved in United States for both subtherapeutic and therapeutic use in poultry under the veterinary feed directive (VDF) - on antimicrobial resistance genes, mobile genetic elements and the functional gene profile of the turkey cecal microbiome.

Antibiotic resistance is a common response of bacteria to antibiotic stress (Gupta et al. 2021), it was therefore not surprising ARG abundance increased immediately after exposure to BMD in both the subtherapeutic and therapeutic groups. However, the continued increase of ARGs only in the therapeutic group and its co-occurrence with MGE even after reducing the BMD dose indicates that the long-term risk of ARGs and their transfer remains a challenge in turkey production after therapeutic use of BMD even after treatment stops. Consistent with our observation of an increase in ARG abundance following the reduction of BMD dose in the therapeutic group, previous studies have reported the persistence of ARGs and antibiotic resistant bacteria even after the withdrawal of the antibiotics to which they were previously exposed (Langlois et al., 1986; Ogunlana et al., 2023).

Interestingly, in our study, therapeutic BMD increased the abundance of ARG multidrug efflux transporters (mdtABC and AcrAB-TolC) and genes conferring resistance to antibiotics other than bacitracin, like tetracycline, beta lactamase, aminoglycoside and vancomycin. Bacitracin binds to the two-component (*baeSR*) system which activates multidrug resistance transporter (*mdtABC*) to pump out bacitracin (Gaetano et al., 2023) and can also increase virulence (Ma et al., 2019). The baeSR – mdtABC interaction was previously shown to increase multidrug resistance in *E. coli* (Nagakubo et al., 2002). Although, most multidrug efflux transporters are encoded on chromosome (Martinez et al., 2009; Paulsen et al., 2001; Saier et al., 1998), AcrAB-TolC has been shown to remove antibiotic from the bacterial cell through active transport, allowing subsequent acquisition of antibiotic-specific resistance genes by plasmid transfer (Nolivos et al., 2019). Taken together, the increase in complete multidrug efflux operons, the co-occurrence of efflux transporters with mobile genetic elements, and the increase in the abundance of resistance gene for antibiotics other than BMD suggests that a therapeutic dose of BMD may increase the risk of multidrug resistance in turkeys by either selecting for multidrug resistant bacteria or by increasing the mobility of ARGs.

We also found that therapeutic BMD administration increased the abundance of T4SS genes, sustained until day 78, and they co-occurred with ARGs. This strengthens evidences that therapeutic BMD facilitates horizontal ARG transfer through conjugation (G. Liu et al., 2019; Dimitriu et al., 2021; Headd & Bradford, 2018), supporting the hypothesis that T4SS play a key role in antibiotic-induced ARG acquisition and dissemination (Bragagnolo et al., 2020; X. Liu et al., 2022; Low et al., 2014). Additionally, prophage genes increased following BMD administration, consistent with reports that antibiotics such as carbadox induce phage gene expression (Johnson et al., 2017). A recent study showed that colibactin, a small genotoxic molecule, produced by several members of the gut microbiome, can induce prophage to lyse their bacterial host as a stress response (Silpe et al., 2022). An important difference with these previous reports is that BMD is not known to cause DNA damage, which is a typical stressor to induce prophage activation. However, high antibiotic concentrations (therapeutic dose) may trigger sustained increase in prophage induction (Wendling et al., 2021). CRISPR-related genes, a bacterial defense system against phage infection, were also increased in the therapeutic group, further indicating that prophage induction may have occurred in these animals. Although there have been contrasting reports regarding the transfer of ARGs by phages through transduction (Torres-Barceló, 2018), the co-occurrence of phage structural proteins with several ARGs in the therapeutic group on day 78 suggests that phage-mediated ARG transfer warrants further investigation. Direct evidence of such transfer could be obtained by isolating and sequencing phage particles directly. Additionally, co-occurrence results should be interpreted with caution because they only provide indirect association but not direct evidence that genes are encoded on the same genetic element. More research is needed to investigate the frequency of T4SS- and prophage-mediated ARG horizontal gene transfer during therapeutic BMD administration. Notable, Barbosa & Levy (2000) proposed a similar mechanism of ARG acquisition via MGEs in livestock under subtherapeutic antibiotic use. However, our findings contrast with their observation, as ARG acquisition through conjugative plasmids predominantly occurred in at therapeutic dose of BMD in turkeys. This hypothesis requires further investigation across varying concentrations of BMD and other antibiotics in common food-producing animals.

Our results showed that BMD treatment, especially the therapeutic dose, significantly decreased the number of ORFs in the turkey cecal microbiome and shifted its functional profile. The dose dependent decrease in ORF diversity suggests that a higher concentration of BMD has a more pronounced impact on the microbiome structure, potentially reducing the community’s functional capabilities. This is consistent with previous studies that have shown that antibiotics can disrupt the microbiome, leading to a decline in functional capacity (Li et al., 2020). Interestingly, despite receiving the same dose as the subtherapeutic group from day 35 onward, the therapeutic group maintained a distinct functional profile, which indicates that therapeutic treatment with BMD might induce a long-lasting shift in microbial function which may potentially alter its development trajectory and resilience to future perturbations (Schwartz et al., 2020).

BMD treatment significantly increased the abundance of genes involved in tryptophan biosynthesis, primarily through the bioconversion of phenylalanine or quinate via chorismite. Previously, metabolomic analysis of these same samples (Johnson et al., 2019) showed that BMD increased tryptophan and its metabolites. This metagenomic study clarifies that BMD administration increased bacterial biosynthesis of tryptophan, suggesting the observed tryptophan increase may be partially attributed to bacterial production. Tryptophan is an important precursor for neurotransmitters like serotonin and neurohormones like melatonin (Parthasarathy et al., 2018) and tryptophan signaling can be influenced by the microbiome (Fu et al., 2022, 2023). Tryptophan metabolites like kynurenine can induce an anti-inflammatory effect (Kanova & Kohout, 2021; Sorgdrager et al., 2019) and increase protein synthesis through the activation of mammalian target of rapamycin (mTOR) (Dukes et al., 2015; Yoon, 2017) which may contribute to animal growth. While the exact role of BMD in modulating tryptophan metabolism remains unclear, our findings suggest a hypothesis that altered tryptophan metabolism is related to the growth promoting effects of BMD in livestock.

Although bacitracin inhibits bacterial growth, it is not clear if it affects all members of the community equally. We hypothesized that the effect of bacitracin is not equal on all members of the community, so we conducted microbial growth rate through the iRep estimates. There was decreased community wide bacterial growth rates in both treatment groups early after BMD exposure on day 7, particularly in the therapeutic group. Over time species, such as *Coprococcus comes* and *Dorea formicigenerans* recovered, suggesting a potential adaptive resilience within the microbiome. A decrease in bacterial growth rate by beta lactam antibiotics was observed in the microbiome of infants which recovered over time (Dubois et al., 2024) similar to what was observed in the BMD groups in our study. Antibiotic growth promoters have been proposed enhance animal growth by reducing microbial load, thereby decreasing immunological energy expenditures and nutrient competition with microbes (Gaskins et al., 2002). This effect of AGP is primarily expected in the small intestine, where most nutrient absorption occurs, rather than in the cecum, as in our study, where microbial fermentation benefits the host (Gaskins, 2000). Since microbial replication rate can be associated with microbial load, BMD could exert selective pressure on the gut microbiota, leading to suppression of specific bacterial populations and potentially influencing animal growth and health over time. Further investigation is needed to elucidate this effect of BMD in both the small intestine and cecum.

## Conclusions

Collectively, this study shows that in-feed BMD, especially at a therapeutic dose, significantly affected the ARG, MGE and metabolic profiles of the turkey cecal microbiome. The enrichment of ARGs and MGEs in response to BMD raises concerns about the potential for increased horizontal gene transfer of ARGs, which may have implications from turkey and human from a one-health perspective. BMD also increased the abundance of tryptophan synthesis genes via the chorismite pathway. Further research should investigate the impact of tryptophan metabolites on host functions. These findings provide insights on risk associated with BMD administration and potential metabolic targets for developing microbiome-modulatory alternatives to AGPs for turkey production and health.

## Methods

### Animal treatment and sample collection

This experiment was approved by the Institutional Animal Care and Use Committee (IACUC) at the National Animal Disease Center (NADC) under protocol ARS-2869. Two hundred and forty male day-of-hatch Nicolas turkey poults (Valley of the Moon Hatchery, Osceola, Iowa) were initially co-housed in a single room for 14 days on a conditioned litter obtained from NADC’s non-antibiotic treated specific-pathogen free Small-Beltsville white turkey flock to homogenize microbiota between birds. These were the same animals as were used in a previous publication (Johnson et al., 2019). Briefly, poults were randomly assigned to three treatment groups: control (no antibiotic), subtherapeutic BMD (50 g/ton feed) and therapeutic BMD (200 g/ton feed). The therapeutic group received 200g/ton BMD for 35 days, followed by 50g/ton until day 78 (Fig. 9). Refer to Johnson et al. (2019) for additional details regarding animal management, diets and housing.

**Figure 9.**
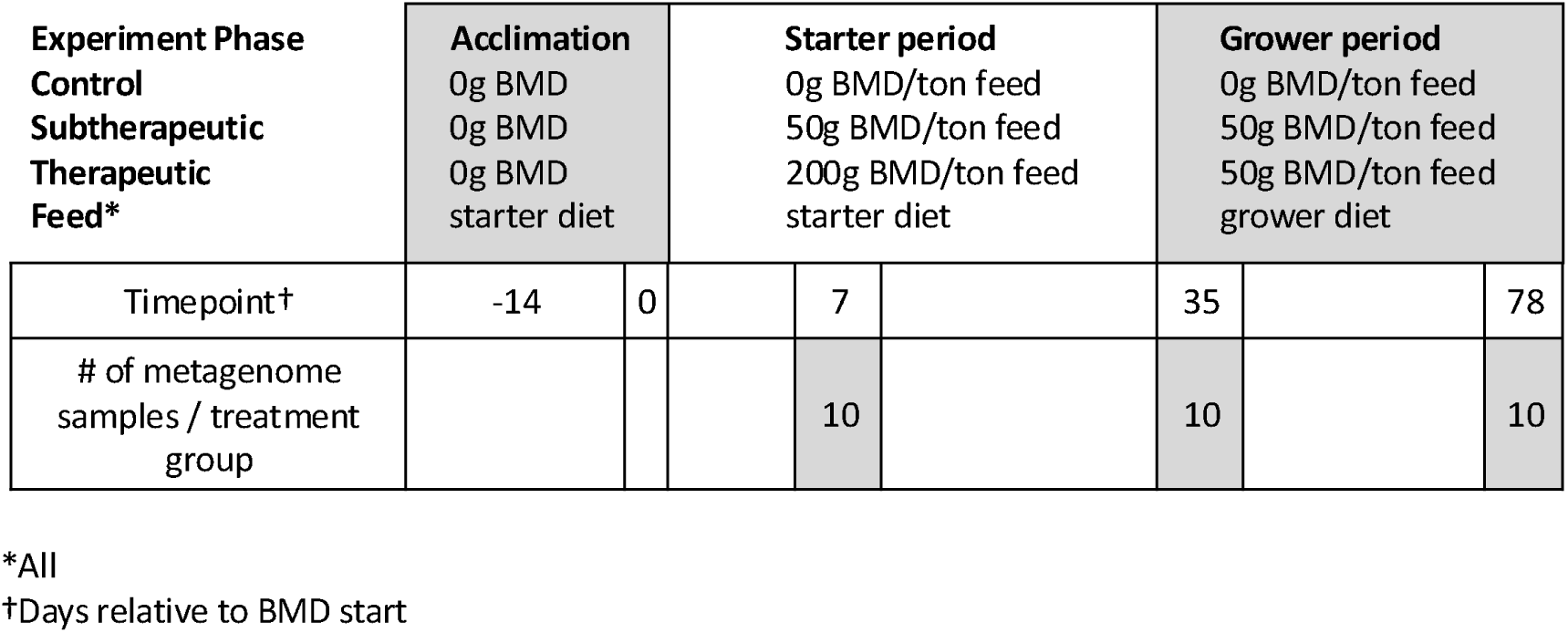
Time sequence (relative to start of BMD treatment) of antibiotic administration, diet and sample collection throughout the study.

### Sample collection and sequencing

Ten turkeys from each treatment group were euthanized after 7, 35 and 78 days of BMD administration. Cecal contents of the euthanized birds were collected for metagenomic analysis. DNA was isolated from the collected cecal samples (PowerMag microbiome DNA/RNA isolation kit, MO-BIO laboratories, Carlsbad, CA). Metagenomic DNA libraries (n = 10 per timepoint/treatment) were generated for short read sequencing using TrueSeq chemistry on an Illumina HiSeq3000 (2 × 150 bp; Iowa State University, Ames, IA) and long read sequencing with the PacBio RSII SMRT sequencing technology (Pacific Biosciences, Menlo Park, CA, USA), in accordance with the manufacturer’s instructions.

### Sequence quality filtering and assembly

Illumina sequences were processed using the BBTools suite (Bushnell et al., 2017) for quality control as follows. For all sequence processing steps, the default parameters were used if no parameters are stated here. Optical duplicates were removed using BBMap (v.38.18) through the clumpify.sh script, while low-quality regions of the flow cell were filtered using filterbytile.sh (removed 2% of reads). Adapter sequences and quality control sequences (PhiX sequences) were removed with BBDuk (v.38.34). Sequences mapping to the turkey genome were then removed with bbsplit.sh. Error correction was performed on the remaining sequences with bbmerge.sh and clumpify.sh before combining them. The combined sequence set was normalized with a target of 100X coverage using bbnorm.sh. One co-assembly was generated from the pooled, normalized Illumina sequences, and the PacBio RSII sequences using metaSPAdes (v.3.12.0, kmer length set at 25, 55, 95, 125) (Nurk et al., 2017). After assembly, contigs shorter than 500 bp were removed. For full details, all files and scripts used in data analysis for this study are available at https://github.com/oluwapaul/BMD-Metagenomics.

### Prediction of open reading frame (ORF) and annotation

Open reading frames (ORFs) in the assembled contigs were predicted with MetaGeneMark (v3.26). The resulting amino acid sequences (.faa) were functionally annotated with the Hidden Markov Model (hmm) - based prediction tool HMMER (version 3.2.1), which queried the TIGRfam database (Release 15.0, downloaded locally) and created a table of hmm-predicted annotations for each gene. The resulting hmm table was converted into a general feature format (.gff) file using an in-house script. The .gff file was used, in combination with previously created .bam files for each treatment group to determine gene counts (Liao et al., 2014). TIGRfam annotation role files (role link and role names) were used to create a combined file of TIGRfam ID, functional annotation, role link, main role, and sub role using an in-house script. To identify ARGs, we used the Resistance Gene Identifier (RGI, version 4.0.3) to compare sequences against the comprehensive antibiotic resistance database (CARD, version 3.2.0; McArthur et al., 2013) with the following parameters: “--input_type contig --low_quality - -local --include_nudge --clean”.

### MAG generation and estimation of replication rates

Assemblies were length-filtered to retain contigs 2: 500 bp, and quality-controlled reads from individual samples were mapped to the co-assembly of each sample using BBMap (v.38.18). The resulting BAM files were processed using the *jgi_summarize_bam_contig_depths* scripts of MetaBAT2 (v.2.12.1; Kang et al., 2019) to calculate within- and between-sample coverages. The mapping files were then used to generate MAGs with MetaBAT2 (-*minContig 1500* and *-maxEdges 200*). MAGs shorter than 5kb were excluded, as recommended in the iRep workflow. CheckM (v.1.0.11; Parks et al., 2015) was used to calculate completeness and contamination of the resulting MAGs. Only MAGs that were 75% complete and had less than 2.5% contamination were retained for replication rate estimation with iRep (Brown et al., 2016). In line with package recommendations, iRep estimates originating from MAGs with lower than 5x coverage were removed. iRep estimates were further filtered to include only those from MAGs that were detected a minimum of four times in all treatment groups per timepoints. A total of 161 MAGs were used for generating 2,122 iRep estimates.

### Network analysis

Network analysis was conducted to identify possible association between ARGs and MGEs (conjugative T4SS, conjugative transposons and phage genes) to further elucidate potential MGEs used for ARG trafficking. The normalized functional gene abundance table was filtered to include only MGEs, which was merged with the normalized ARG abundance table. Co-occurrence analysis was performed with the *rcorr* function from the corrplot package in R using Spearman correlation with Benjamin-Hochberg procedure for the correction of false discovery. Pairs of genes with strong correlation coefficients (rho > 0.8) that were statistically significant (q < 0.05) were retained in the analysis. The co-occurrence network was visualized with Cytoscape (v3.9.0; Shannon et al., 2003).

### Statistical Analysis

All statistical analysis was done in R v. 4.2.2.(R Core Team, 2014). Functional genes were normalized to the total number of sequences per million reads (reads per million, rpm) while ARGs were normalized to the total number of sequences and gene length per million reads (reads per kilobase per million reads mapped, rpkm). The effects of BMD on functional gene and ARG profiles were examined with a non-metric multidimensional scaling (NMDS) based on Bray– Curtis dissimilarity distance, calculated with normalized ORF and ARG count tables using the *vegdist* function from the vegan package in R. PERMANOVA with the Holms correction was used to detect statistical differences in community composition between treatment groups. Kruskal-Wallis test was used to test significant statistical differences in the alpha diversity of functional genes and ARG between the different treatment groups. Differential abundance analyses of both the functional and ARG was done with the DESeq2 package. To explore the associations between time, treatment and community wide bacterial growth rates, linear mixed models implemented by the R package lme4 (Bates et al., 2015) were employed. A model with the formula iRep ∼ day * treatment + (1|genome) was used. P-value for fixed effect of BMD treatment were generated using the lmerTest package using Satterthwaite’s approximation (Kuznetsova et al., 2020). Contrasts of interest were extracted with the emmeans package (Lenth, 2022) and adjusted P-values were generated (using FDR control) Significance was set at P-value < 0.05. Data wrangling and plotting were accomplished with tidyverse (Wickham et al., 2019), phyloseq and ggplot2 packages.

## Availability of data and materials

All raw sequencing reads are available in the NCBI sequence read archive (SRA) under the project accession PRJNA988023. All additional files and scripts used in data analysis for this study are available at https://github.com/oluwapaul/BMD-Metagenomics.

## Abbreviations

AGP: Antibiotic growth promoter
ANOVA: Analysis of variance
ARG: Antibiotic resistant gene
BMD: Bacitracin methylene disalicylate
CARD: Comprehensive Antibiotic Resistance Database
DAG: Differentially abundant gene
FDA: Food and Drug Administration
iRep: Index of replication
MAG: metagenome-assembled genomes
MGE: mobile genetic elements
NMDS: Non-metric multidimensional scaling
ORF: Open reading frame
PERMANOVA: Permutational Multivariable Analysis of Variance
RGI: Resistance Gene Identifier
T4SS: Type IV secretion system
VFD: Veterinary feed directive.

## Acknowledgement

We thank the staff at USDA-ARS National Animal Disease Center, Ames, Iowa for assistance in carrying out the animal study.

## Authors’ contributions

T.L. and T.A.J. conceived the study, T.L., and T.A.J. collected the samples, P.O., C.L.W., J.T., and T.A.J performed the data analysis, P.O., C.L.W., J.T., T.L. and T.A.J discussed the results, P.O. wrote the manuscript, T.L. and T.A.J. edited the manuscript. All authors read and approved the final manuscript.

## Corresponding author

Correspondence to Timothy A. Johnson

## Declarations

### Ethics approval and consent to participate

This experiment was approved by the Institutional Animal Care and Use Committee (IACUC) at the National Animal Disease Center (NADC) and the experiment was carried out according to the approved protocol ARS-2869.

## Consent for publication

Not applicable

## Supplementary Information

**Additional file 1:** List of differentially abundant genes in BMD treatment groups relative to time-matched control. **Table S1**. Subtherapeutic relative to control on day 7. **Table S2.** Therapeutic relative to control on day 7. **Table S3.** Subtherapeutic relative to control on day 35. **Table S4.** Therapeutic relative to control on day 35. **Table S5**. Subtherapeutic relative to control on day 78. **Table S6.** Therapeutic relative to control on day 78.

**Additional file 2:** List of differentially abundant ARGs in BMD treatment groups relative to time-matched controls. **Table S7**. Subtherapeutic relative to control on day 7. **Table S8.** Therapeutic relative to control on day 7. **Table S9.** Therapeutic relative to Subtherapeutic on day 7. **Table S10.** Subtherapeutic relative to control on day 35. **Table S11.** Therapeutic relative to control on day 35. **Table S12.** Therapeutic relative to Subtherapeutic on day 35. **Table S13**. Subtherapeutic relative to control on day 78. **Table S14.** Therapeutic relative to control on day 78. **Table S15.** Therapeutic relative to Subtherapeutic on day 78.

**Additional file 3:** Summary of ARG-MGE correlation network parameters (**Table S16**).

**Additional file 4:** Supplemental Figures. **Figure S1.** Differentially abundant genes from DESeq2 in treatment groups relative to control on day 7 for A. Subtherapeutic and B. Therapeutic. **Figure S2.** Differentially abundant genes from DESeq2 in treatment groups relative to control on day 35 for A. Subtherapeutic and B. Therapeutic. **Figure S3.** Differentially abundant genes from DESeq2 in treatment groups relative to control on day 78 for A. Subtherapeutic and B. Therapeutic. **Figure S4.** Differentially abundant genes of the type IV secretion system (T4SS) relative to control on day 35 in A. Subtherapeutic, B. Therapeutic, and differentially abundant phage genes on day 35 in C. Subtherapeutic and D. Therapeutic. **Figure S5.** Differentially abundant conjugative transposons in Subtherapeutic relative to control on day 7 A, day 35 B, and day 78 C. Differentially abundant conjugative transposons in Therapeutic relative to control on day 7 D, day 35 E and day 78 F. **Figure S6.** Network analysis showing the spearman correlation between ARG and MGE in turkey cecal microbiota on day 7. **Figure S7.** Network analysis showing the spearman correlation between ARG and MGE in turkey cecal microbiota on day 35. **Figure S8.** Network analysis showing the spearman correlation between ARG and phage-related genes in turkey cecal microbiota on day 7. **Figure. S9.** Network analysis showing the spearman correlation between ARG and phage-related genes in turkey cecal microbiota on day 35. **Figure S10.** Network analysis showing the spearman correlation between ARG and phage-related genes in turkey cecal microbiota on day 78. **Figure S11.** Abundance of genes involved in tryptophan synthesis from either phenylalanine or quinate. **Figure S12.** Effect of BMD administration on community wide replication rates overtime.

