## Additional file 4 for "In-feed bacitracin methylene disalicylate alters microbiota function and increases antibiotic resistance in a dose-dependent manner"

**Figure S8.** Network analysis showing the spearman correlation between ARG and phage-related genes in turkey cecal microbiota on day 7. Pairwise correlation of padj < 0.05 and rho > 0.8 was considered. The nodes represent the ARG and MGE, the size of the node represent its number of connections (Degree). A-C represent ARG– phage gene on day 7 in control subtherapeutic and therapeutic group respectively. Triangles are phage genes while circle are ARG. Color of the circle indicate different ARG mechanism, green is antibiotic efflux, blue is antibiotic inactivation, red is antibiotic target alteration, purple is antibiotic target protection while light green is antibiotic target replacement.

**Figure S9.** Network analysis showing the spearman correlation between ARG and phage-related genes in turkey cecal microbiota on day 35. Pairwise correlation of padj < 0.05 and rho > 0.8 was considered. The nodes represent the ARG and MGE, the size of the node represent its number of connections (Degree). A-C represent ARG– phage gene on day 35 in control subtherapeutic and therapeutic group. Triangles are phage gens while circle are ARG. Color of the circle indicate different ARG mechanism, green is antibiotic efflux, blue is antibiotic inactivation, red is antibiotic target alteration, purple is antibiotic target protection while light green is antibiotic target replacement.

**Figure S10.** Network analysis showing the spearman correlation between ARG and phage-related genes in turkey cecal microbiota on day 78. Pairwise correlation of padj < 0.05 and rho > 0.8 was considered. The nodes represent the ARG and MGE, the size of the node represent its number of connections (Degree). A-C represent ARG– phage gene on day 35 in control subtherapeutic and therapeutic group. Triangles are phage gens while circle are ARG. Color of the circle indicate different ARG mechanism, green is antibiotic efflux, blue is antibiotic inactivation, red is antibiotic target alteration, purple is antibiotic target protection while light green is antibiotic target replacement.

**Figure S11.** Abundance of genes involved in tryptophan synthesis from either phenylalanine or quinate. A. Aminodeoxychorismate lyase B. 3-Phosphoshikimate-1-carboxyvinyltransferase C. 3-deoxy-7phosphoheptulonate synthase D. membrane bound PQQ dependent dehydrogenase glucose quinate shikimate family E. Chorismate synthase F. Isochorismate synthase G. Indolepyruvate ferredoxin oxidoreductase alpha subunit H. Indolepyruvate ferredoxin oxidoreductase beta subunit I. Phenylacetate CoA oxygenase PaaG subunit J. Phenylacetate CoA oxygenase PaaH subunit K. Phenylacetate CoA oxygenase PaaI subunit L. Phenylacetate CoA oxygenase PaaJ subunit M. Phenylacetate CoA oxygenase reductase PaaK subunit N. Phenylacetate CoA ligase.

**Figure S12.** Effect of BMD administration on community wide replication rates overtime. Estimated iRep replication mean on day 7 (A), day 35 (B), and day 78 (C). Estimated iRep replication rates in representative MAGs on day 7 (D), day 35 (E), and day 78 (F).

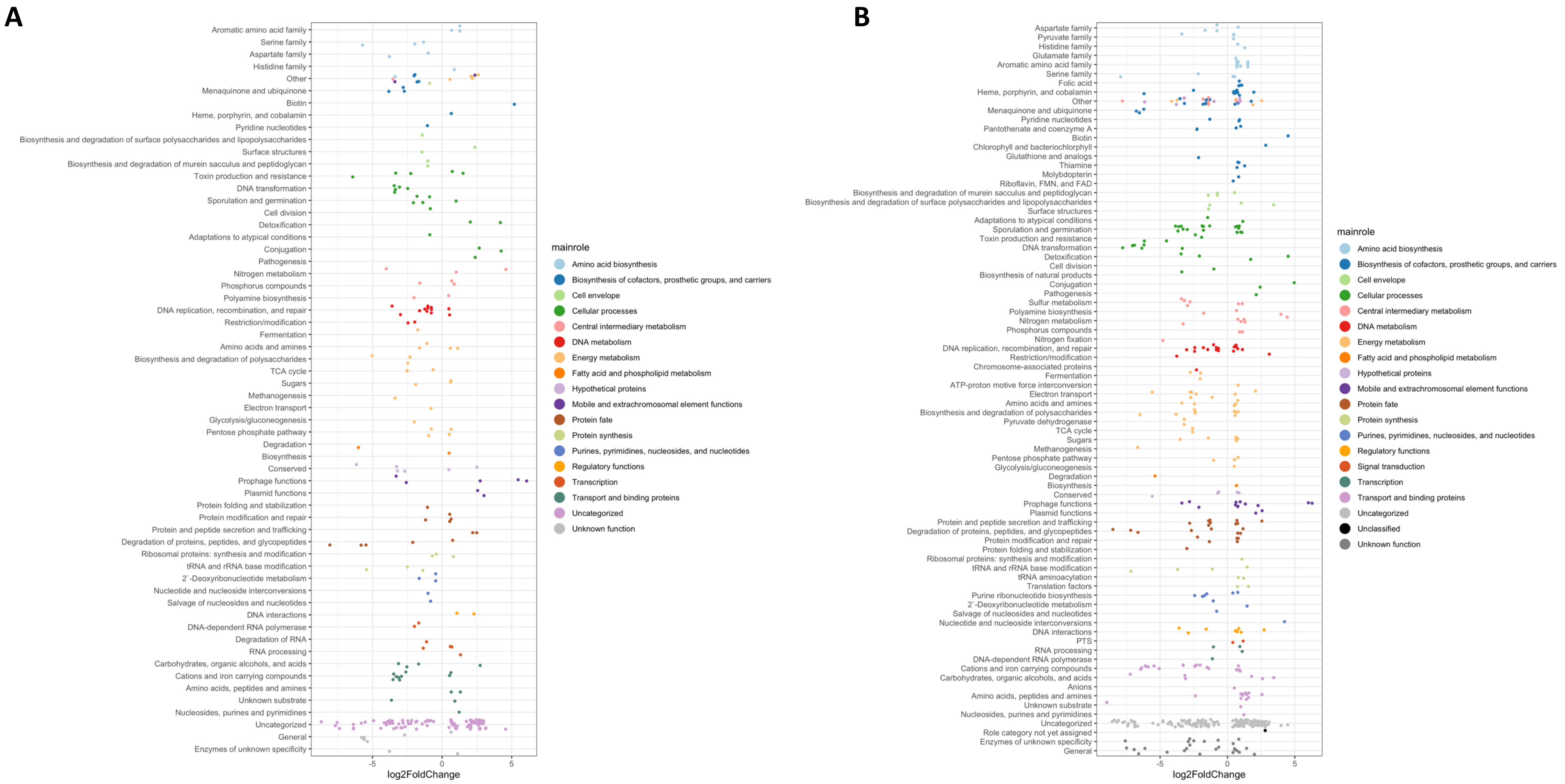


**Figure S1.** Differentially abundant genes from DESeq2 in treatment groups relative to control on day 7 for A. Subtherapeutic and B. Therapeutic.

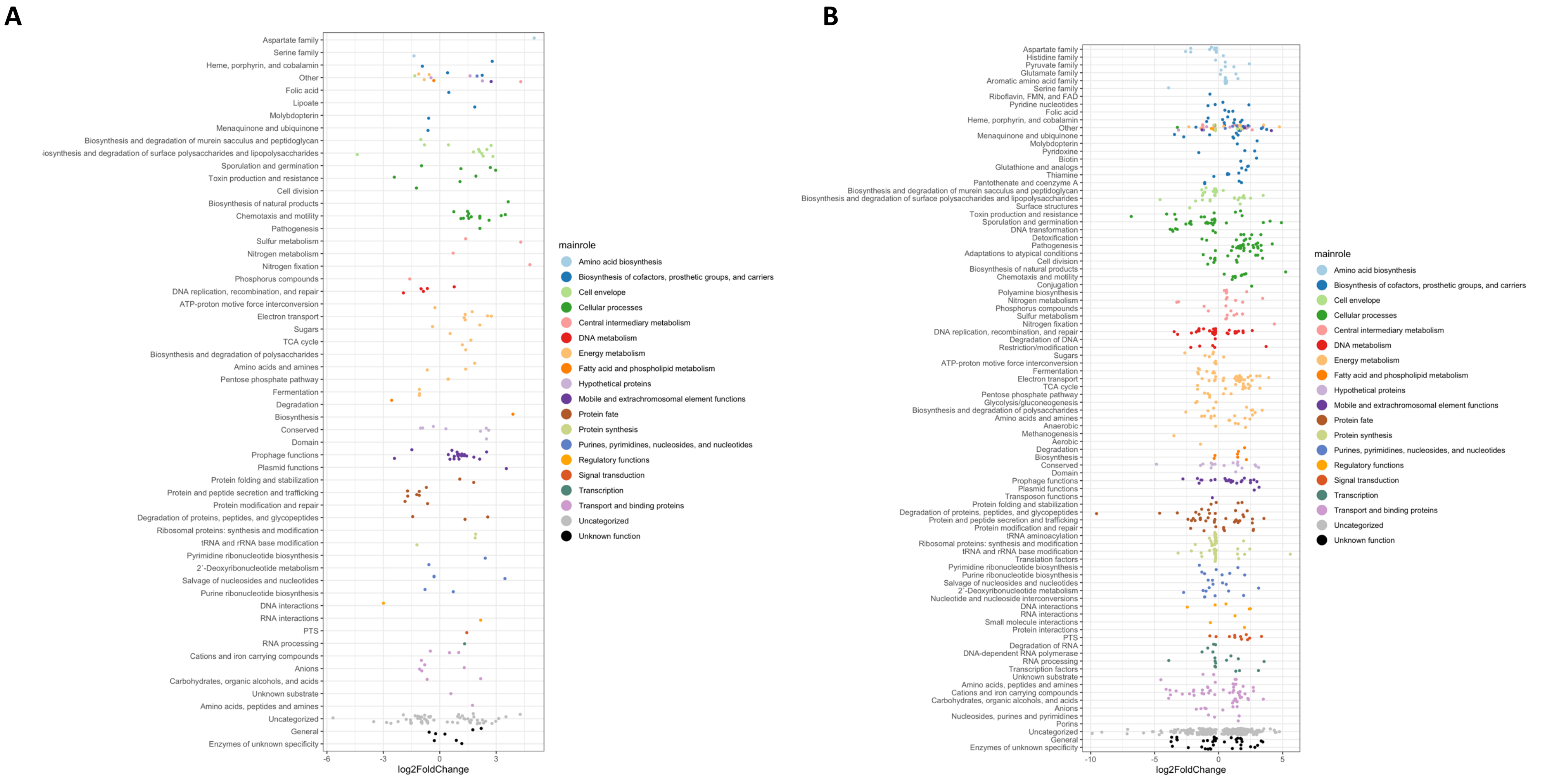


**Figure S2.** Differentially abundant genes from DESeq2 in treatment groups relative to control on day 35 for A. Subtherapeutic and B. Therapeutic.

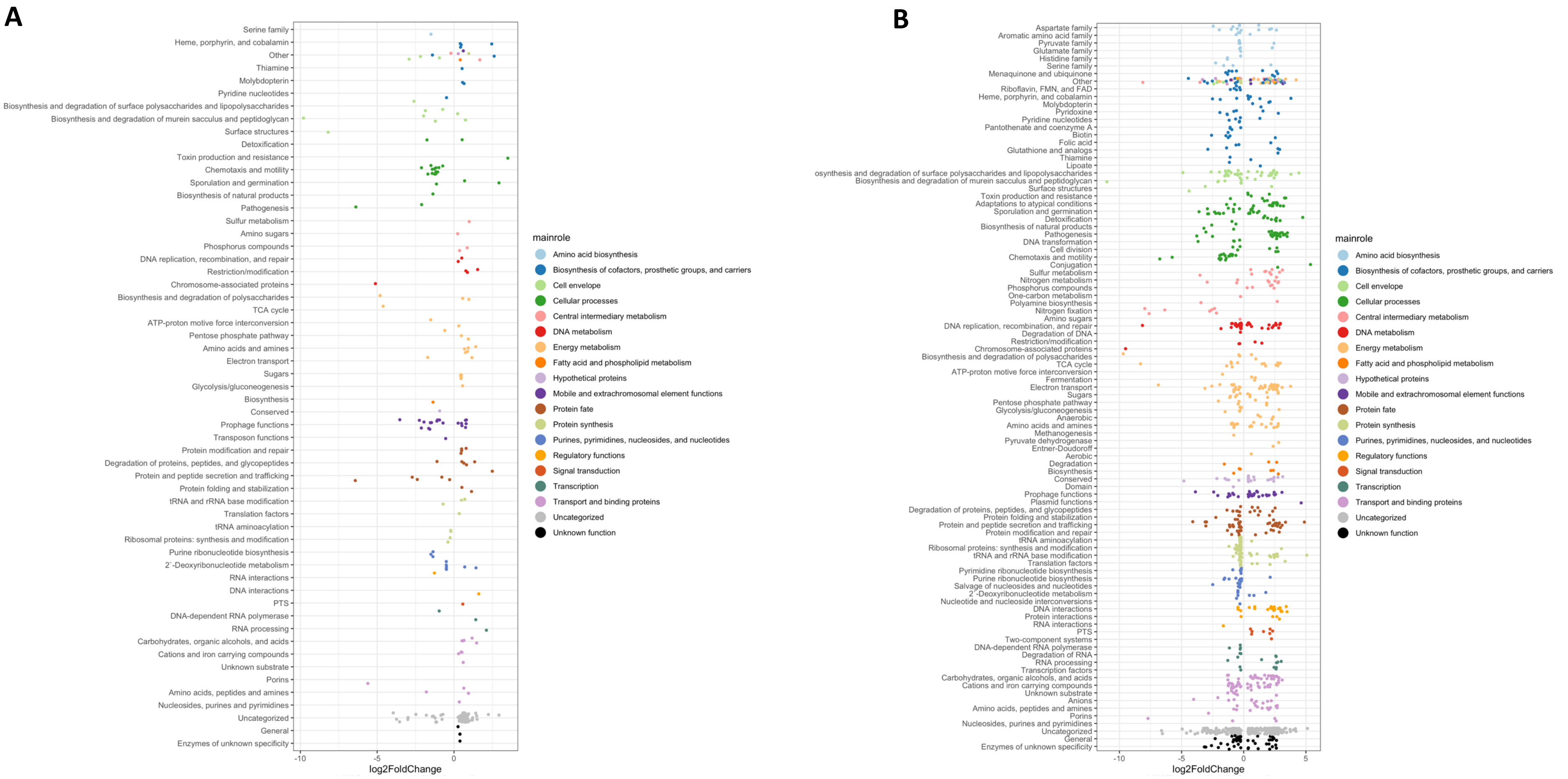


**Figure S3.** Differentially abundant genes from DESeq2 in treatment groups relative to control on day 78 for A. Subtherapeutic and B. Therapeutic.

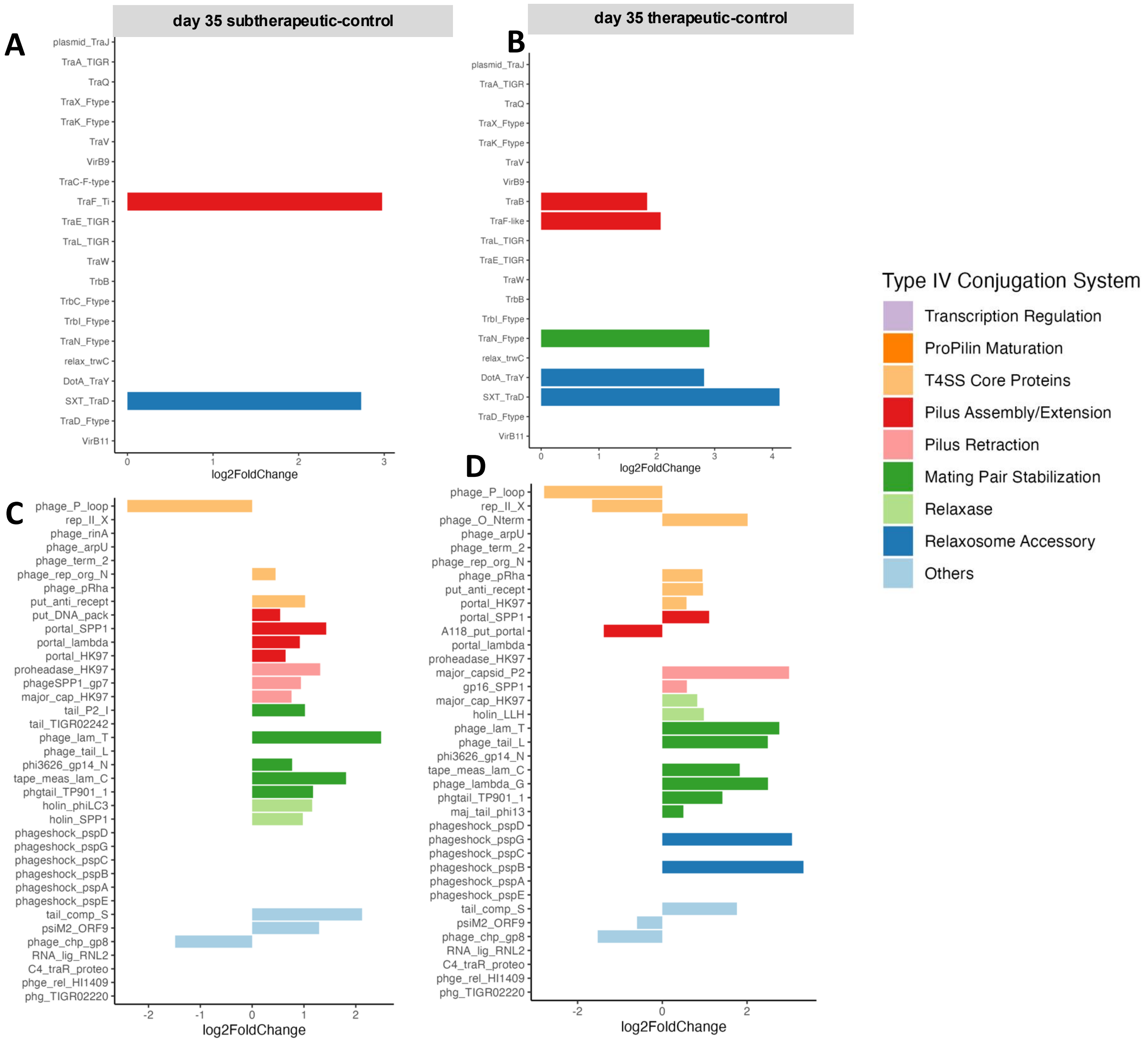


**Figure S4.** Differentially abundant genes of the type IV secretion system (T4SS) relative to control on day 35 in A. Subtherapeutic, B. Therapeutic, and differentially abundant phage genes on day 35 in C. Subtherapeutic and D. Therapeutic.

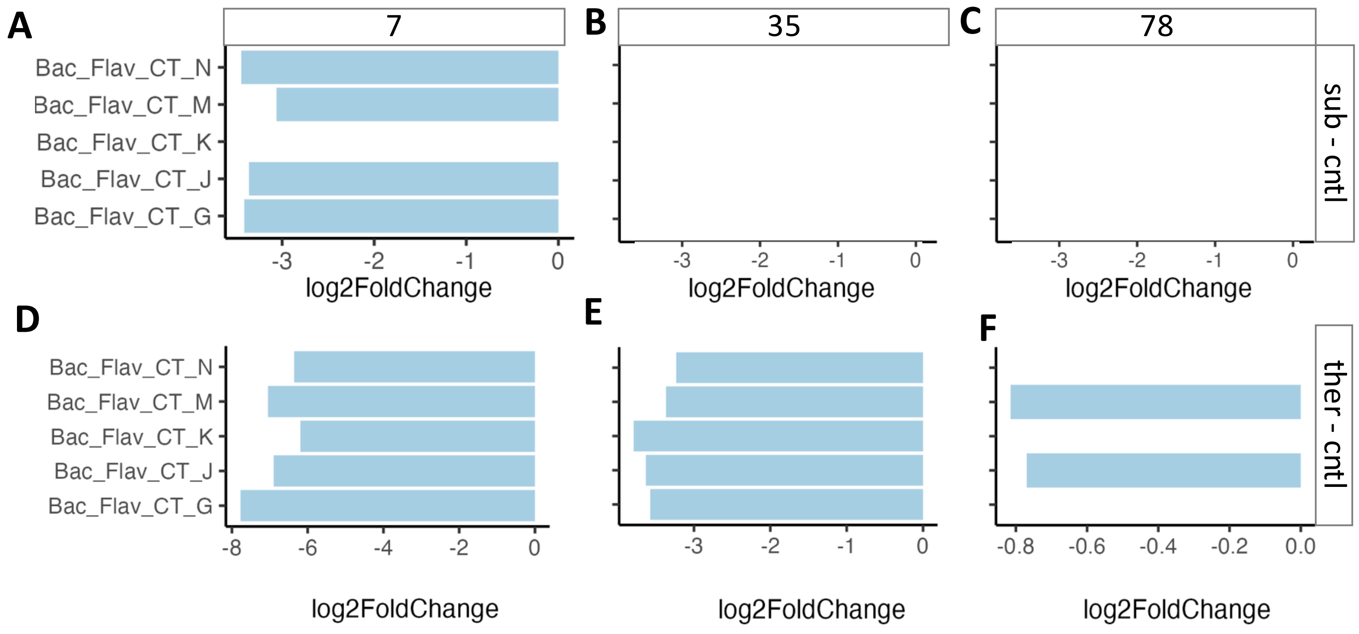


**Figure S5.** Differentially abundant conjugative transposons in Subtherapeutic relative to control on day 7 A, day 35 B, and day 78 C. Differentially abundant conjugative transposons in Therapeutic relative to control on day 7 D, day 35 E and day 78 F.

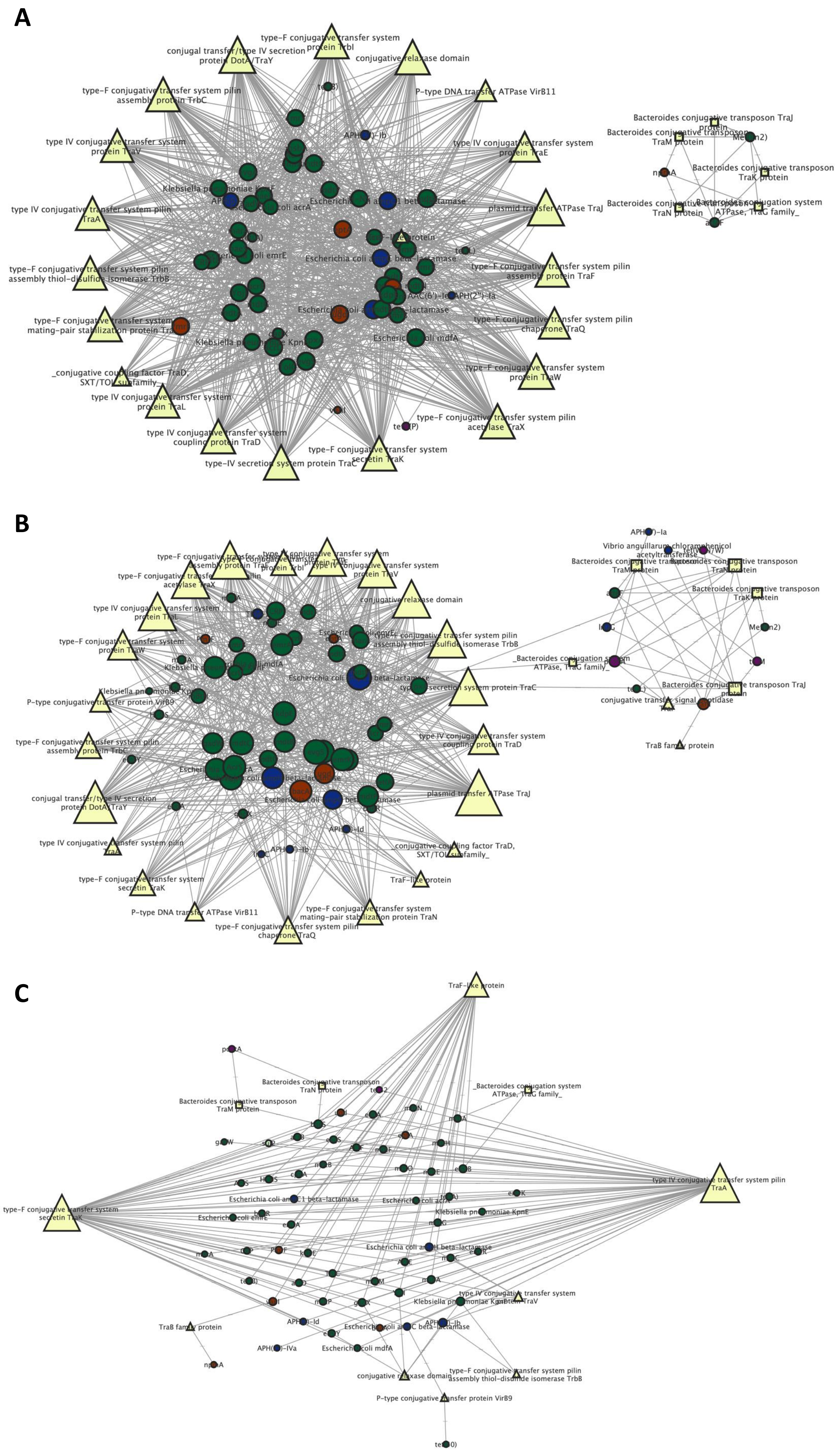


**Figure S6.** Network analysis showing the spearman correlation between ARG and MGE in turkey cecal microbiota on day 7. Pairwise correlation of padj < 0.05 and rho > 0.8 was considered. The nodes represent the ARG and MGE, the size of the node represent its number of connections (Degree). A-C represent ARG– conjugation gene (both T4SS and conjugative transposon) on day 7 in control subtherapeutic and therapeutic group respectively. Color of the circle indicate different ARG mechanism, green is antibiotic efflux, blue is antibiotic inactivation, red is antibiotic target alteration, purple is antibiotic target protection while light green is antibiotic target replacement.

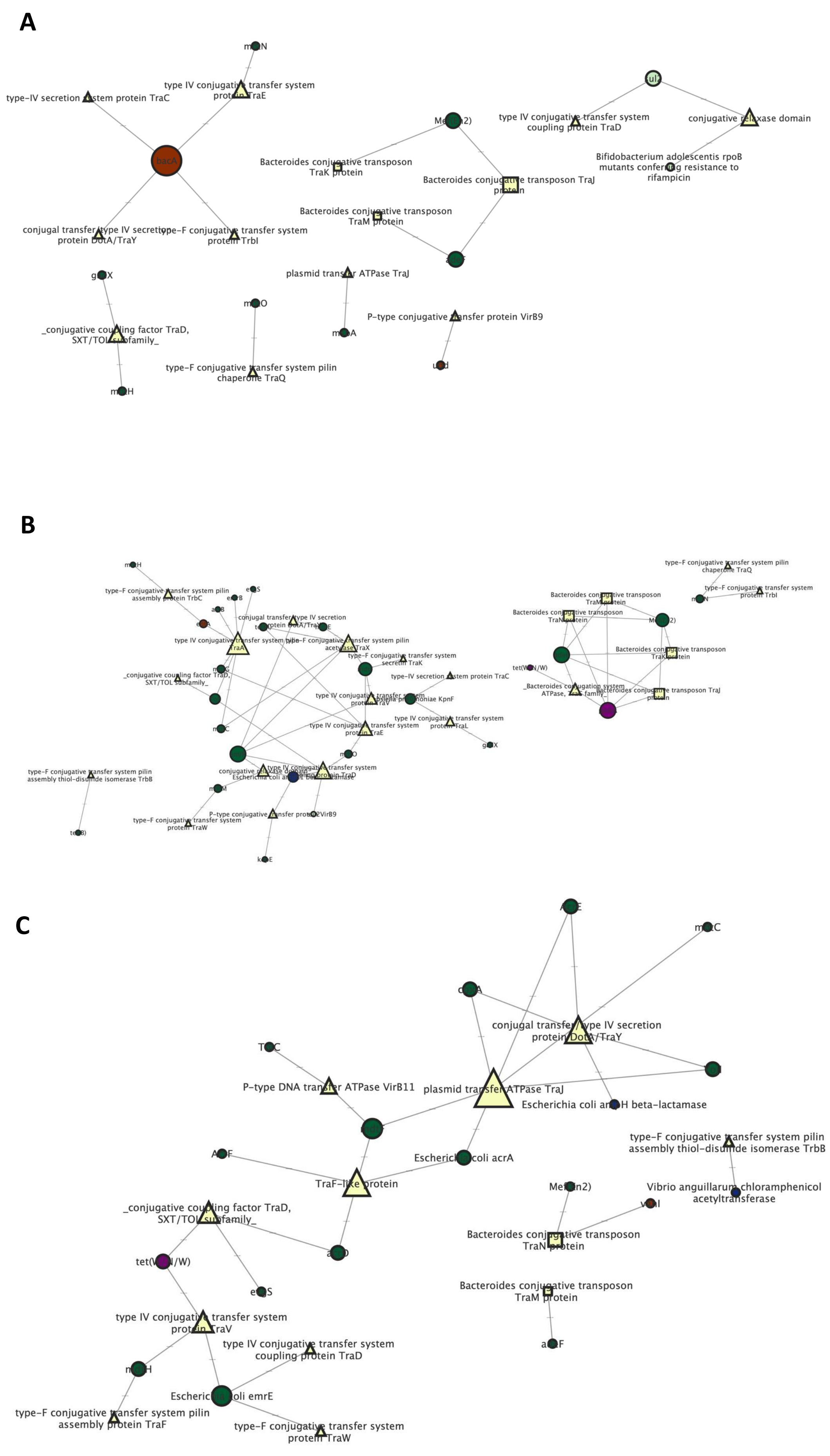


**Figure S7.** Network analysis showing the spearman correlation between ARG and MGE in turkey cecal microbiota on day 35. Pairwise correlation of padj < 0.05 and rho > 0.8 was considered. The nodes represent the ARG and MGE, the size of the node represent its number of connections (Degree). A-C represent ARG– conjugation gene (both T4SS and conjugative transposon) on day 35 in control subtherapeutic and therapeutic group. Triangles are T4SS, square are conjugative transposon while circle are ARG. Color of the circle indicate different ARG mechanism, green is antibiotic efflux, blue is antibiotic inactivation, red is antibiotic target alteration, purple is antibiotic target protection while light green is antibiotic target replacement.

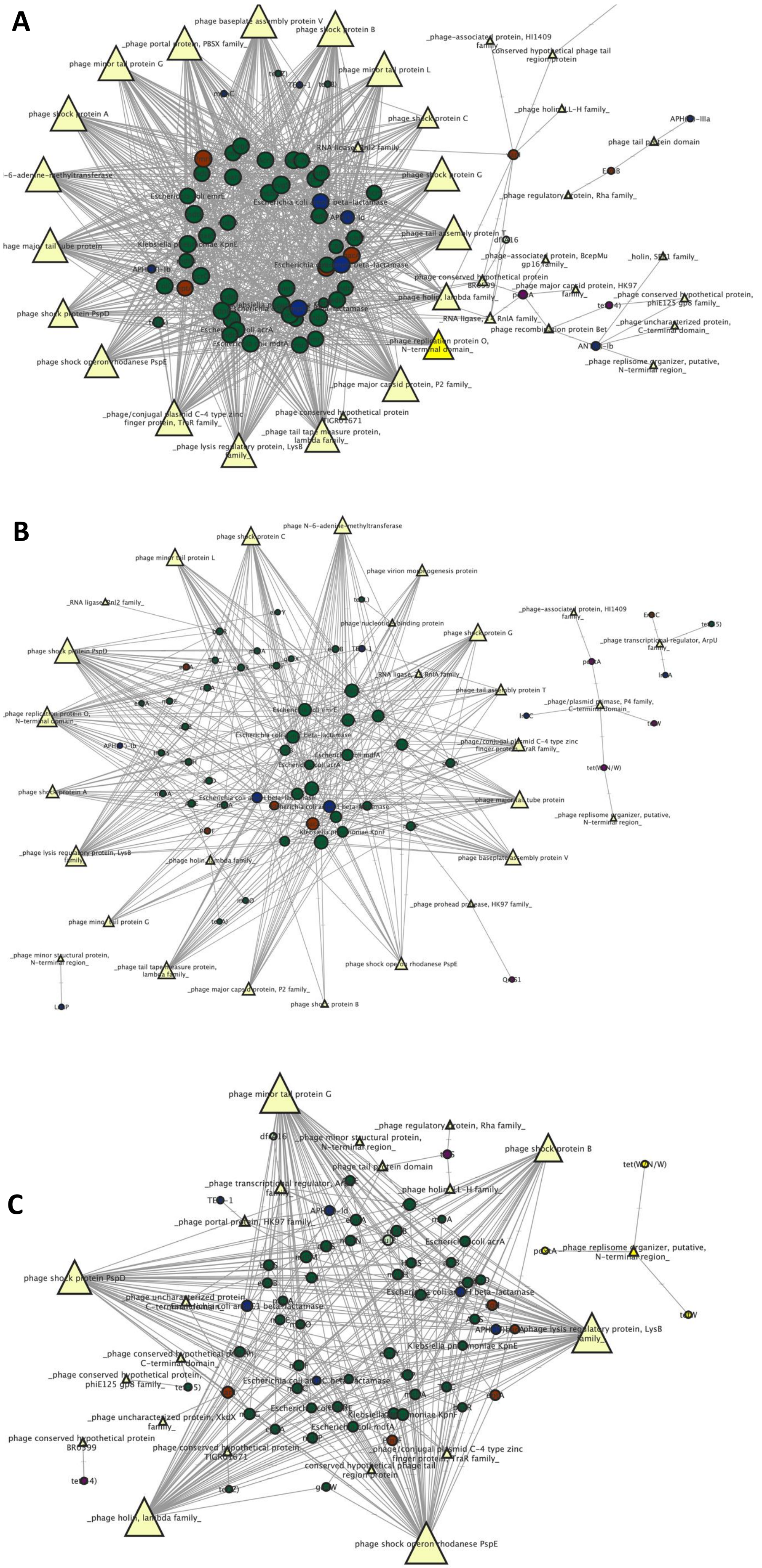


**Figure S8.** Network analysis showing the spearman correlation between ARG and phage-related genes in turkey cecal microbiota on day 7. Pairwise correlation of padj < 0.05 and rho > 0.8 was considered. The nodes represent the ARG and MGE, the size of the node represent its number of connections (Degree). A-C represent ARG– phage gene on day 7 in control subtherapeutic and therapeutic group respectively. Triangles are phage genes while circle are ARG. Color of the circle indicate different ARG mechanism, green is antibiotic efflux, blue is antibiotic inactivation, red is antibiotic target alteration, purple is antibiotic target protection while light green is antibiotic target replacement.

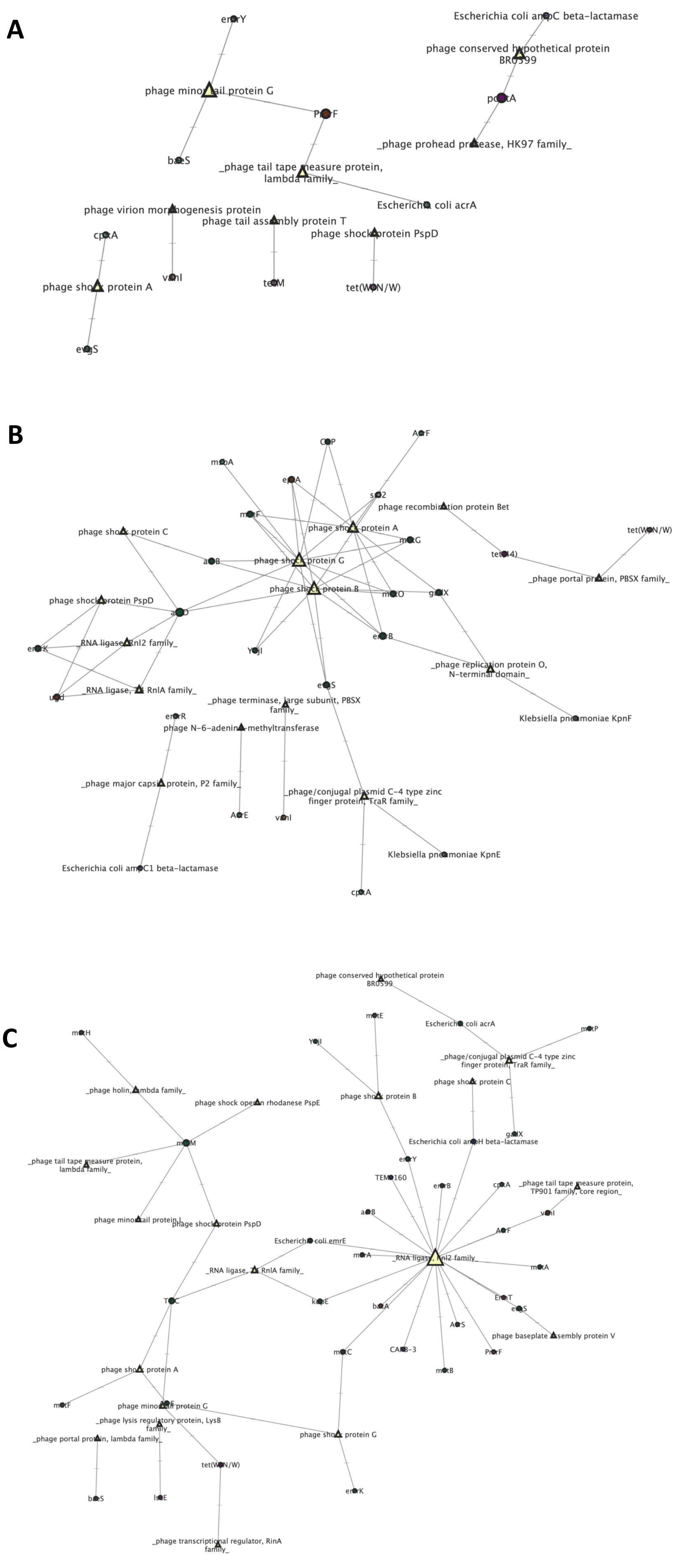


**Figure S9.** Network analysis showing the spearman correlation between ARG and phage-related genes in turkey cecal microbiota on day 35. Pairwise correlation of padj < 0.05 and rho > 0.8 was considered. The nodes represent the ARG and MGE, the size of the node represent its number of connections (Degree). A-C represent ARG– phage gene on day 35 in control subtherapeutic and therapeutic group. Triangles are phage gens while circle are ARG. Color of the circle indicate different ARG mechanism, green is antibiotic efflux, blue is antibiotic inactivation, red is antibiotic target alteration, purple is antibiotic target protection while light green is antibiotic target replacement.

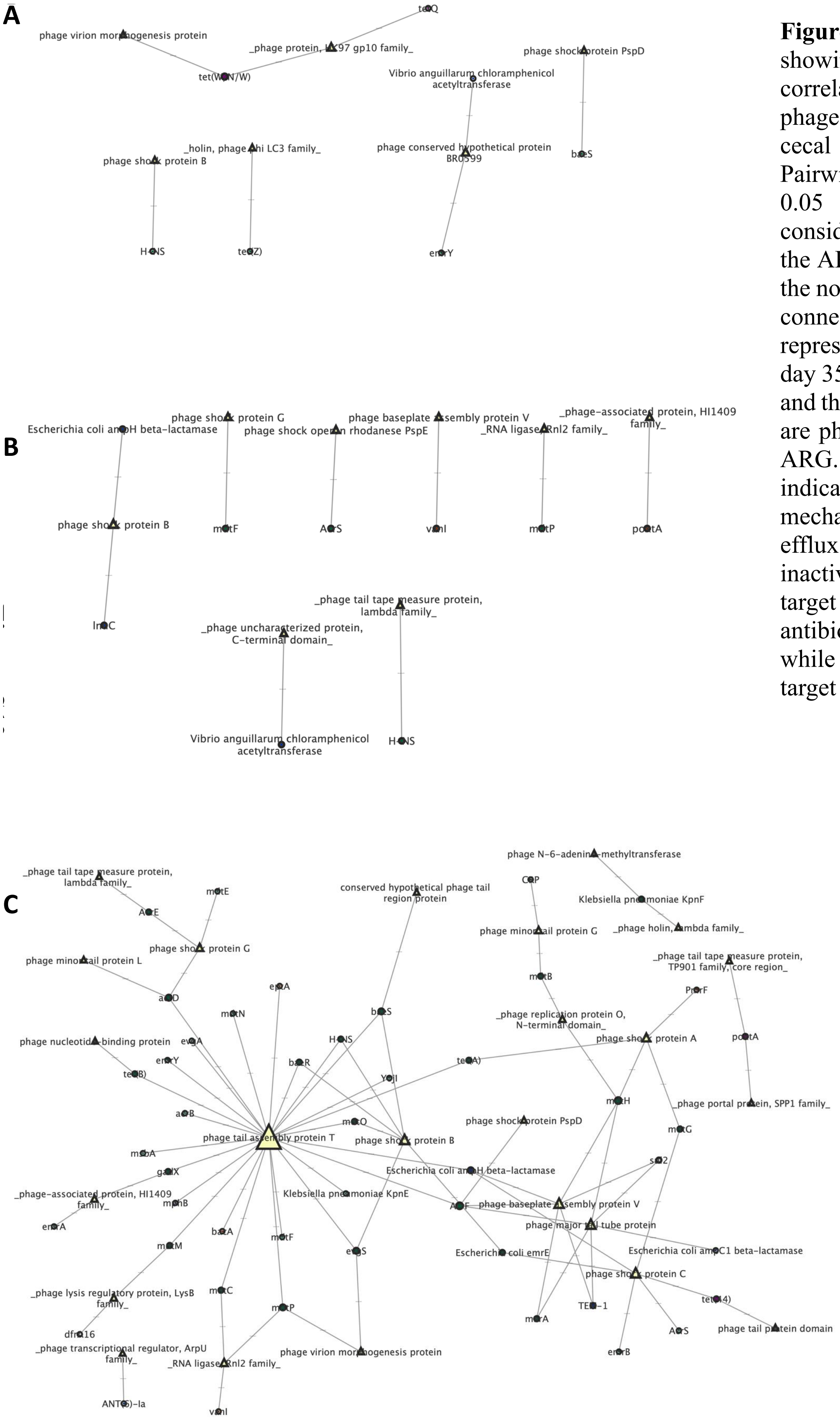


**Figure S10.** Network analysis showing the spearman correlation between ARG and phage-related genes in turkey cecal microbiota on day 78. Pairwise correlation of padj < 0.05 and rho > 0.8 was considered. The nodes represent the ARG and MGE, the size of the node represent its number of connections (Degree). A-C represent ARG– phage gene on day 35 in control subtherapeutic and therapeutic group. Triangles are phage gens while circle are ARG. Color of the circle indicate different ARG mechanism, green is antibiotic efflux, blue is antibiotic inactivation, red is antibiotic target alteration, purple is antibiotic target protection while light green is antibiotic target replacement.

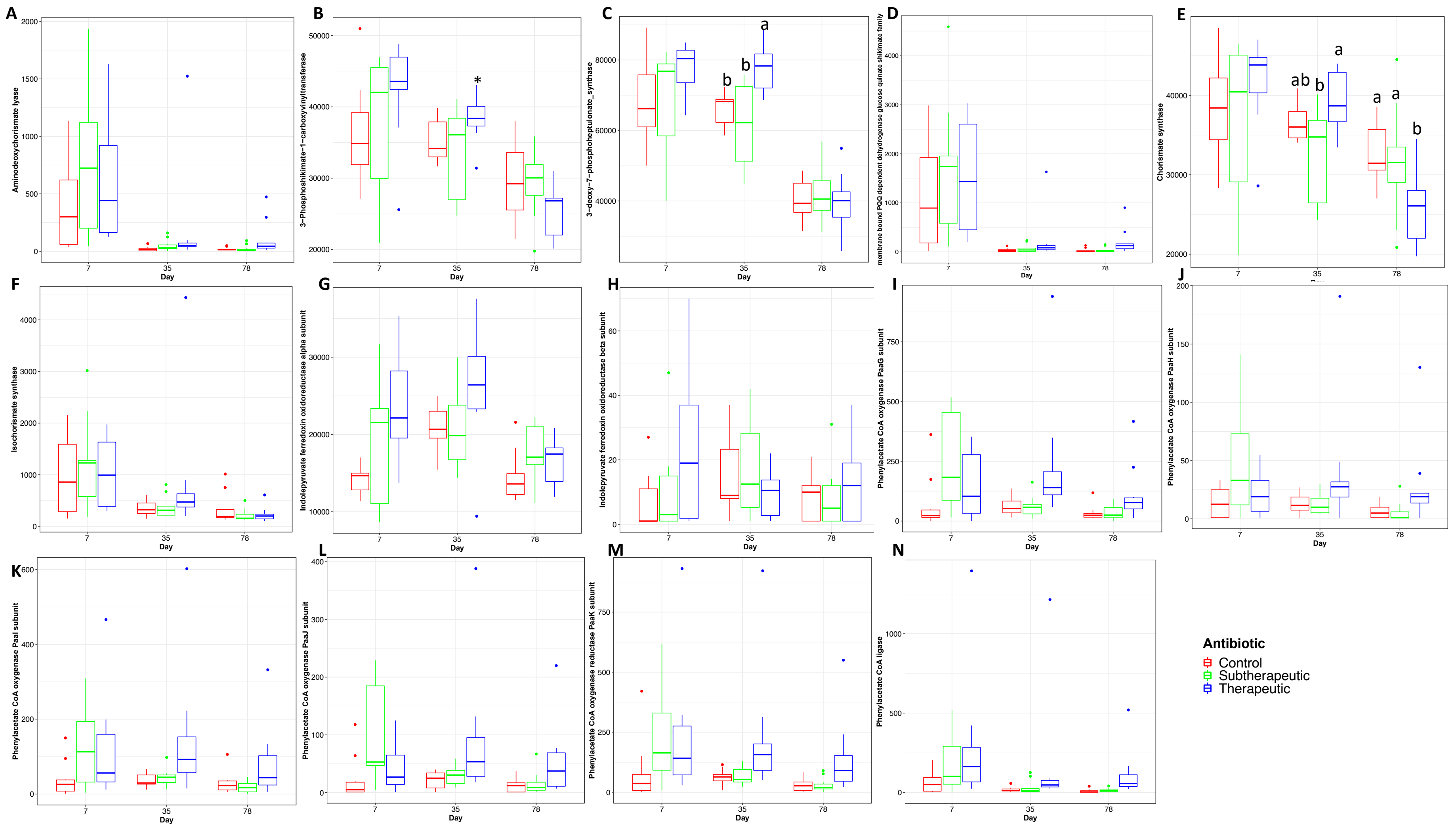

**Figure S11.** Abundance of genes involved in tryptophan synthesis from either phenylalanine or quinate. A. Aminodeoxychorismate lyase B. 3-Phosphoshikimate-1-carboxyvinyltransferase C. 3-deoxy-7phosphoheptulonate synthase D. membrane bound PQQ dependent dehydrogenase glucose quinate shikimate family E. Chorismate synthase F. Isochorismate synthase G. Indolepyruvate ferredoxin oxidoreductase alpha subunit H. Indolepyruvate ferredoxin oxidoreductase beta subunit I. Phenylacetate CoA oxygenase PaaG subunit J. Phenylacetate CoA oxygenase PaaH subunit K. Phenylacetate CoA oxygenase PaaI subunit L. Phenylacetate CoA oxygenase PaaJ subunit M. Phenylacetate CoA oxygenase reductase PaaK subunit N. Phenylacetate CoA ligase.

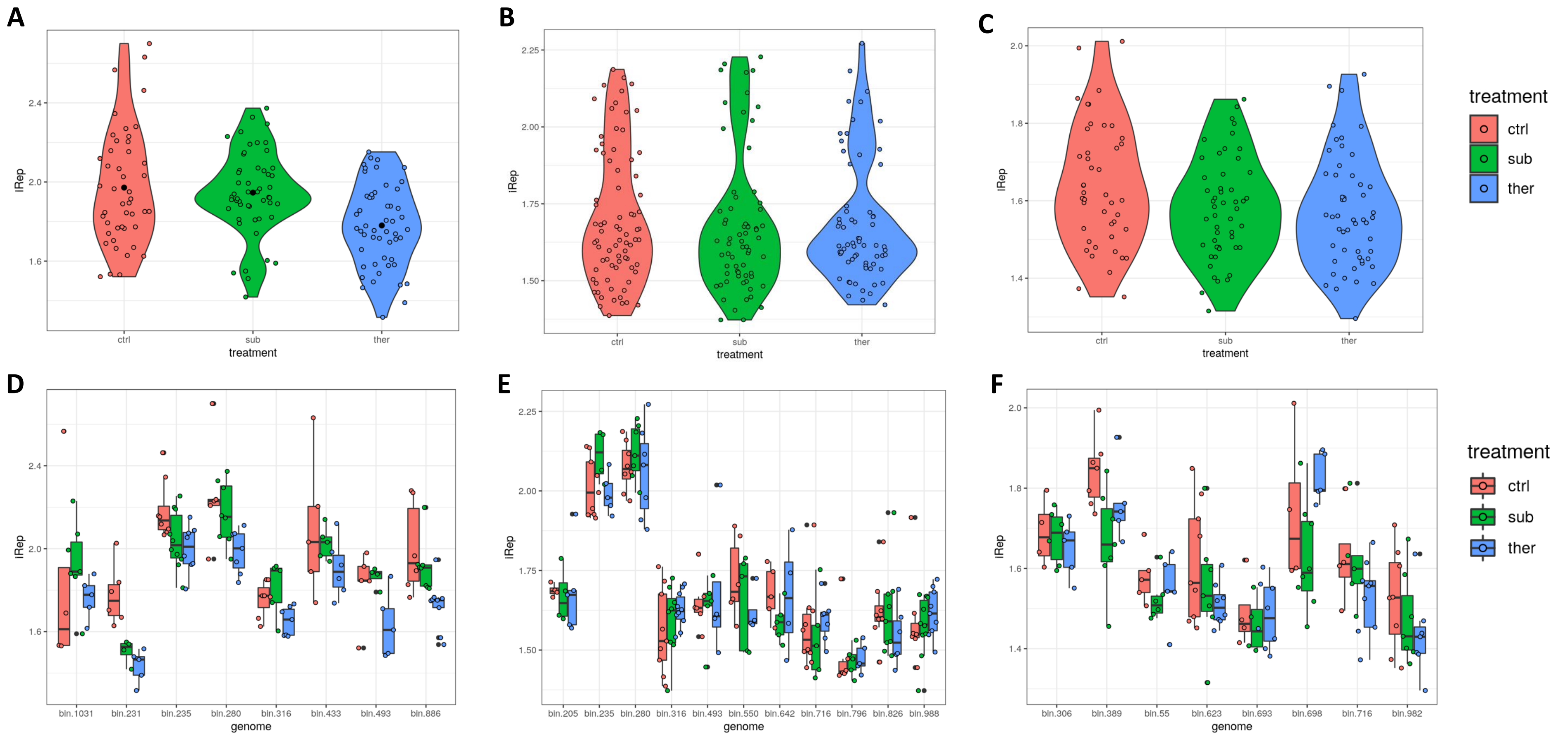


**Figure S12.** Effect of BMD administration on community wide replication rates overtime. Estimated iRep replication mean on day 7 (A), day 35 (B), and day 78 (C). Estimated iRep replication rates in representative MAGs on day 7 (D), day 35 (E), and day 78 (F).
